# Bcr4 is a Chaperone for the Inner Rod Protein in the *Bordetella* Type III Secretion System

**DOI:** 10.1101/2021.09.28.462275

**Authors:** Masataka Goto, Akio Abe, Tomoko Hanawa, Masato Suzuki, Asaomi Kuwae

## Abstract

*Bordetella bronchiseptica* injects virulence proteins called effectors into host cells via a type III secretion system (T3SS) conserved among many Gram-negative bacteria. Small proteins called chaperones are required for stabilizing some T3SS components or localizing them to the T3SS machinery. In a previous study, we identified a chaperone-like protein named Bcr4 that regulates T3SS activity in *B*. *bronchiseptica*. Bcr4 does not show strong sequence similarity to well-studied T3SS proteins of other bacteria, and its function remains to be elucidated. Here, we investigated the mechanism by which Bcr4 controls T3SS activity. A pull-down assay revealed that Bcr4 interacts with BscI, based on its homology to other bacterial proteins, to be an inner rod protein of the T3SS machinery. An additional pull-down assay using truncated Bcr4 derivatives and secretion profiles of *B*. *bronchiseptica* producing truncated Bcr4 derivatives showed that the Bcr4 C-terminal region is necessary for the interaction with BscI and activation of the T3SS. Moreover, the deletion of BscI abolished the secretion of type III secreted proteins from *B*. *bronchiseptica* and the translocation of a cytotoxic effector into cultured mammalian cells. Finally, we showed that BscI is unstable in the absence of Bcr4. These results suggest that Bcr4 supports the construction of the T3SS machinery by stabilizing BscI. This is the first demonstration of a chaperone for the T3SS inner rod protein among the virulence bacteria possessing the T3SS.

**Importance:** The type III secretion system (T3SS) is a needle-like complex that projects outward from bacterial cells. *Bordetella bronchiseptica* uses the T3SS to inject virulence proteins into host cells. Our previous study reported that a protein named Bcr4 is essential for the secretion of virulence proteins from *B. bronchiseptica* bacterial cells and delivery through the T3SS. Because other bacteria lack a Bcr4 homologue, the function of Bcr4 has not been elucidated. In this study, we discovered that Bcr4 interacts with BscI, a component of the T3SS machinery. We showed that a *B. bronchiseptica* BscI-deficient strain was unable to secrete type III secreted proteins. Furthermore, in a *B. bronchiseptica* strain that overproduces T3SS component proteins, Bcr4 is required to maintain BscI in bacterial cells. These results suggest that Bcr4 stabilizes BscI to allow construction of the T3SS in *B. bronchiseptica*.

## Introduction

The genus *Bordetella* consists of Gram-negative bacteria that infect the respiratory tracts of mammals including humans. *Bordetella pertussis* causes a severe coughing attack called whooping cough in humans (1, 2). *Bordetella bronchiseptica* causes atrophic rhinitis in pigs and kennel cough in dogs (1, 2). These *Bordetella* spp. harbor a virulence factor secretion apparatus called the type III secretion system (T3SS).

The T3SS consists of a basal body that penetrates the inner and outer membranes of the bacteria and a needle structure that protrudes outside the bacteria, and is conserved in many Gram-negative bacteria such as *Yersinia*, *Salmonella*, and *Pseudomonas*. *B. bronchiseptica* uses the T3SS to inject virulence factors, called effectors, into host cells to disrupt the physiological functions of host cells (3). Once the basal body and the export apparatus of the T3SS are completed, the type III secreted proteins are secreted in a fixed order. First, the components (Table S1) of the needle structure (SctF) and the inner rod (SctI) that ties the needle structure to the basal body are secreted. After the needle is completed, translocators (SctE and SctB) are secreted, which form small pores in the host cell membrane to create a pathway for effectors. Finally, the effectors translocate into the host cell via the T3SS (4, 5). In *B. bronchiseptica*, BopB (SctE) (6), BopD (SctB) (7), and Bsp22 (SctA) (8) function as translocators (Table S1), while BteA (also referred to as BopC) (9, 10), BopN (11, 12), and BspR (also referred to as BtrA) (13, 14) function as effectors. Bsp22 is located at the tip of the needle and bridges the needle to the pore-forming factors embedded in the plasma membrane (15). BteA has been shown to cause membrane-disrupting cytotoxicity in mammalian cells (10). BspR is a regulator that represses transcription of the *bteA* gene and genes on the *bsc* locus (Fig. S1), where genes encoding type III secreted proteins and components of T3SS are located (14, 16). According to secondary structure prediction—e.g., the predicted positions of helix, and the overall structure of the operon—the BscF and BscI of *B. bronchiseptica* correspond to *Yersinia* needle YscF (SctF) and inner rod YscI (SctI), respectively (Table S1).

In addition, many type III secreted proteins have unique chaperones that are involved in stabilizing the substrate and preventing premature polymerization of the substrate in the bacterial cytosol, and then in efficiently transporting the substrate to the T3SS machinery (17–20). For example, PscE and PscG function as chaperones of the needle PscF (SctF) in *Pseudomonas* (Table S1). These chaperones stabilize PscF in the bacterial cytosol and are involved in PscF secretion through the T3SS (21). In bacteria such as *Yersinia* and *Pseudomonas*, an inner rod chaperone is thought to exist, because the inner rod is secreted out of the bacterial cell and polymerized (22, 23). However, a chaperone for the inner rod has not been reported.

So far, a chaperone-like protein called Bcr4 has been identified in *B. bronchiseptica* (24). *Bordetella* Bcr4 is highly conserved among *B. pertussis*, *B. parapertussis*, and *B. bronchiseptica* (Fig. S2). In this study, we attempted to identify factors that interact with Bcr4 in order to investigate how Bcr4 is involved in the T3SS regulation.

## Results

### Bcr4 binds to BscI, an inner rod protein of the *Bordetella* type III secretion system

The results of a previous study suggested that Bcr4 is a chaperone for components of the type III secretion system (T3SS) (24). T3SS chaperones are known to be involved in substrate stability and efficient transport of substrates to the T3SS machinery (17, 19, 20). In addition, it is generally known that the genes of these chaperones are localized adjacent to genes encoding their substrates (25). On the *B. bronchiseptica* S798 chromosome, the genes encoding BcrH2, BscI, BscJ, and BscK are located in the vicinity of the *bcr4* gene (Fig. 1A), and these proteins are predicted to function as a translocator chaperone, inner rod (SctI), inner membrane ring (SctJ), and ATPase cofactor (SctK), respectively (Table S1) (5, 26). To test whether Bcr4 binds to BscI, BscJ or BscK, we added *E. coli* lysates containing the V5-tagged target proteins (BscI-V5, BscJ-V5 or BscK-V5) to Strep-Tactin beads loaded with the purified Strep-tagged Bcr4 (Bcr4-Strep), and then performed the pull-down assay. The supernatant fraction (Sup) and pellet fraction (Pellet) samples were prepared, separated by SDS-PAGE, and subjected to Western blotting with anti-V5 antibody (Fig. 1B). When the beads were washed with TBS, the V5 signal of the BscI-V5 pellet sample was detected in the beads loaded with Bcr4, but not in the unloaded beads (Fig. 1B). The V5 signals of the BscK-V5 and BscJ-V5 pellet samples were detected in both the Bcr4-loaded beads and the unloaded beads when washed with TBS, and not detected when washed with TBS containing 0.1% Triton X-100 (Fig. 1B). These results suggest that Bcr4 binds to BscI, an inner rod protein of the *Bordetella* T3SS.

**Fig. 1.**
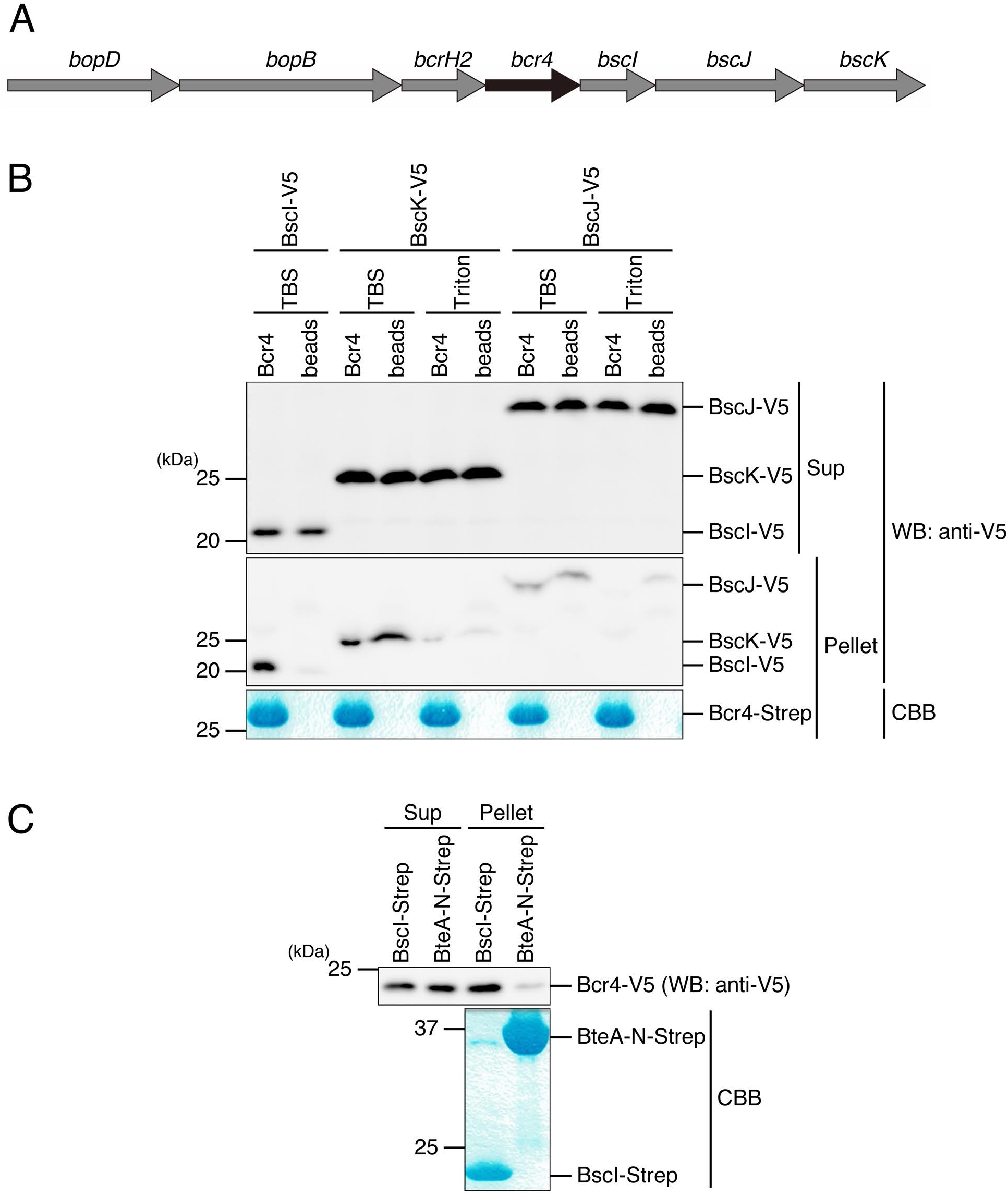
The interaction of Bcr4 with inner rod protein BscI. (A) The *bcr4* gene and its peripheral genes localized in the T3SS apparatus locus (*bsc* locus) on the *B. bronchiseptica* S798 chromosome are depicted. (B) The purified Bcr4 (Bcr4-Strep) was loaded on Strep-Tactin beads. Then, the Bcr4-Strep-loaded beads (Bcr4) and the beads alone (beads) were mixed with the lysates prepared from *E. coli* BL21 producing BscI, BscK or BscJ tagged with V5 (BscI-V5, BscK-V5 or BscJ-V5), respectively. After 3 h incubation at 4°C, each supernatant was prepared as the supernatant fraction (Sup) sample, and each pellet was washed with TBS or TBS containing 0.1% Triton X-100 (Triton) and prepared as the pellet fraction (Pellet) sample. The Sup and Pellet samples were separated by SDS-PAGE and analyzed by Western blotting (WB) with anti-V5 antibody (top). Pellet samples were also stained with Coomassie Brilliant Blue (CBB, bottom). (C) The purified BscI (BscI-Strep) or N-terminal moiety of BteA (amino acids region 1–312, BteA-N-Strep) were loaded on Strep-Tactin beads. Then, the beads were mixed with the lysate prepared from *E. coli* BL21 to produce Bcr4 tagged with V5 (Bcr4-V5) at 4°C for 3 h. The prepared Sup and Pellet samples were separated by SDS-PAGE and analyzed by WB with anti-V5 antibody (top). Pellet samples were also stained with CBB (bottom). Experiments were performed at least three times, and representative data are shown.

Next, to confirm that Bcr4 binds to BscI, a pull-down assay was performed by adding *E. coli* lysate containing V5-tagged Bcr4 (Bcr4-V5) to Strep-Tactin beads with the purified Strep-tagged BscI (BscI-Strep) or BteA N-terminal 1–312 amino acids region (BteA-N-Strep). BteA is a protein secreted from the type III secretion system and interacts with its cognate chaperone BtcA through the N-terminal (9). As a result, the V5 signal was detected in the pellet sample of the BscI-Strep, but was hardly detected in that of the BteA-N-Strep (Fig. 1C). These results strongly suggested that Bcr4 binds to BscI.

### The C-terminal region of Bcr4 is required for the binding of Bcr4 to BscI

Next, to investigate the Bcr4 region responsible for the binding to BscI, we produced Strep-tagged full-length Bcr4 (Bcr4-FL-Strep), Bcr4 lacking amino acids region 58–109 (Bcr4Δ58-109-Strep), and Bcr4 lacking amino acids region 110–173 (Bcr4Δ110-173-Strep) in *E. coli* (Fig. 2A). A pull-down assay was performed by adding *E. coli* lysate containing V5-tagged BscI (BscI-V5) to Strep-Tactin beads with the purified Bcr4-FL-Strep, Bcr4Δ58-109-Strep, or Bcr4Δ110-173-Strep. The Sup and Pellet samples were then prepared, separated by SDS-PAGE, and subjected to Western blotting using anti-V5 antibody (Fig. 2B). As a result, the V5 signal was detected in the pellet sample of Bcr4-FL-Strep, but not in those of Bcr4Δ58-109-Strep and Bcr4Δ110-173-Strep. Although it is still unknown which region of Bcr4 directly interacts with BscI, our results strongly suggest that both the Bcr4-58-109 and Bcr4-110-173 regions are required for the interaction.

**Fig. 2.**
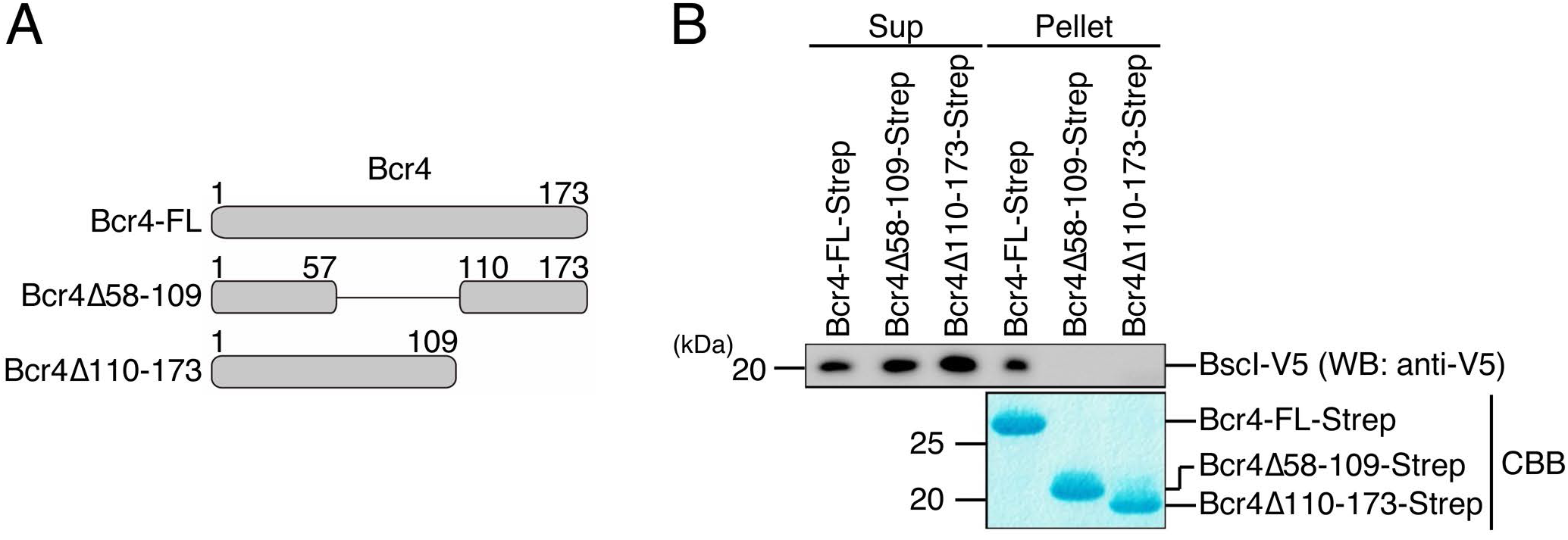
Pull-down assays between BscI and truncated versions of Bcr4. (A) Bcr4 derivatives used for pull-down assay are depicted. (B) The purified Bcr4-FL, Bcr4Δ58-109 or Bcr4Δ110-173 (Bcr4-FL-Strep, Bcr4Δ58-109-Strep or Bcr4Δ110-17-Strep) was loaded on Strep-Tactin beads, and then the beads were mixed with the lysate prepared from *E. coli* BL21, producing BscI-V5, respectively. After 3 h incubation at 4°C, the supernatant fraction (Sup) and pellet fraction (Pellet) samples were prepared. The Sup and Pellet samples were separated by SDS-PAGE and analyzed by Western blotting (WB) with anti-V5 antibody (top), and stained with Coomassie Brilliant Blue (CBB, bottom). Experiments were performed at least three times, and representative data are shown.

### The C-terminal region of Bcr4 is required for T3SS activity

The results in Figure 2 suggest that the C-terminal region of Bcr4 is required for binding to BscI. Therefore, we introduced plasmids encoding the full-length FLAG-tagged Bcr4 (Bcr4-FL-FLAG), Bcr4 with the 5 C-terminal amino acids deleted (Bcr4Δ169-173-FLAG), Bcr4 with the 10 C-terminal amino acids deleted (Bcr4Δ164-173-FLAG), and Bcr4 with the 15 C-terminal amino acids deleted (Bcr4Δ159-173-FLAG) (Fig. 3 A) into a Bcr4-deficient strain (Δ*bcr4*) (Δ*bcr4*+*bcr4*-FL-FLAG, Δ*bcr4*+*bcr4*Δ169-173-FLAG, Δ*bcr4*+*bcr4*Δ164-173-FLAG and Δ*bcr4*+*bcr4*Δ159-173-FLAG, respectively). Whole cell lysates (WCL) and culture supernatant fraction (CS) samples were prepared from these strains and separated by SDS-PAGE. Western blottings were then carried out using antibodies against Bcr4, FLAG, BteA (an effector and a type III secreted protein), BopD (SctB, a translocator and a type III secreted protein) or RpoB (an internal control of WCL) (Fig. 3B). Since the antibody against Bcr4 was generated using its 18 C-terminal amino acids (amino acids (aa) 156–173) as the antigen peptide, we considered that it may not recognize the Bcr4 partial deletion mutant proteins used in this experiment. Therefore, we also performed Western blotting using anti-FLAG antibody (Fig. 3B). As a result, when anti-Bcr4 antibody was used against the WCL samples, signals were detected in the wild-type, Δ*bcr4*+*bcr4*-FL-FLAG and Δ*bcr4*+*bcr4* Δ169-173-FLAG, but almost no signals were detected in Δ*bcr4*+*bcr4*Δ164-173-FLAG and Δ*bcr4*+*bcr4*Δ159-173-FLAG (Fig. 3B). On the other hand, when anti-FLAG antibody was used, signals were detected in Δ*bcr4*+*bcr4*Δ164-173-FLAG and Δ*bcr4*+*bcr4*Δ159-173-FLAG (Fig. 3B). These results confirmed that the Bcr4 partial deletion mutant proteins were produced in the Δ*bcr4* strain. When anti-BteA or anti-BopD antibodies were used against the CS samples, signals were detected in the wild-type, Δ*bcr4*+*bcr4*-FL-FLAG, Δ*bcr4*+*bcr4*Δ169-173-FLAG and Δ*bcr4*+*bcr4*Δ164-173-FLAG, respectively, but not in Δ*bcr4* or Δ*bcr4*+*bcr4*Δ159-173-FLAG (Fig. 3B). We attempted to create *B. bronchiseptica* strains that produce shorter Bcr4, e.g. amino acid regions 1-57, 58-109, and 110-173, however, those truncated Bcr4 were produced at very low levels (data not shown). Therefore, we were unable to evaluate whether or not these truncated proteins were functional in *B. bronchiseptica*. We then infected L2 cells (a rat lung epithelial cell line) with the wild-type, Δ*bcr4*, Δ*bcr4*+*bcr4*-FL-FLAG, Δ*bcr4*+*bcr4*Δ169-173-FLAG, Δ*bcr4*+*bcr4*Δ164-173-FLAG or Δ*bcr4*+*bcr4*Δ159-173-FLAG strain at a multiplicity of infection (MOI) of 50 for 1 h and measured the amounts of LDH released into the culture medium as an index of cytotoxicity. As a result, LDH was detected in the medium of cells infected with Δ*bcr4*+*bcr4*-FL-FLAG, Δ*bcr4*+*bcr4*Δ169-173-FLAG or Δ*bcr4*+*bcr4*Δ164-173-FLAG, but not in the medium of cells infected with Δ*bcr4*+*bcr4*Δ159-173-FLAG (Fig. 3C). These results suggest that the region required for the T3SS function is located in the 159–163 amino acids region of Bcr4.

**Fig. 3.**
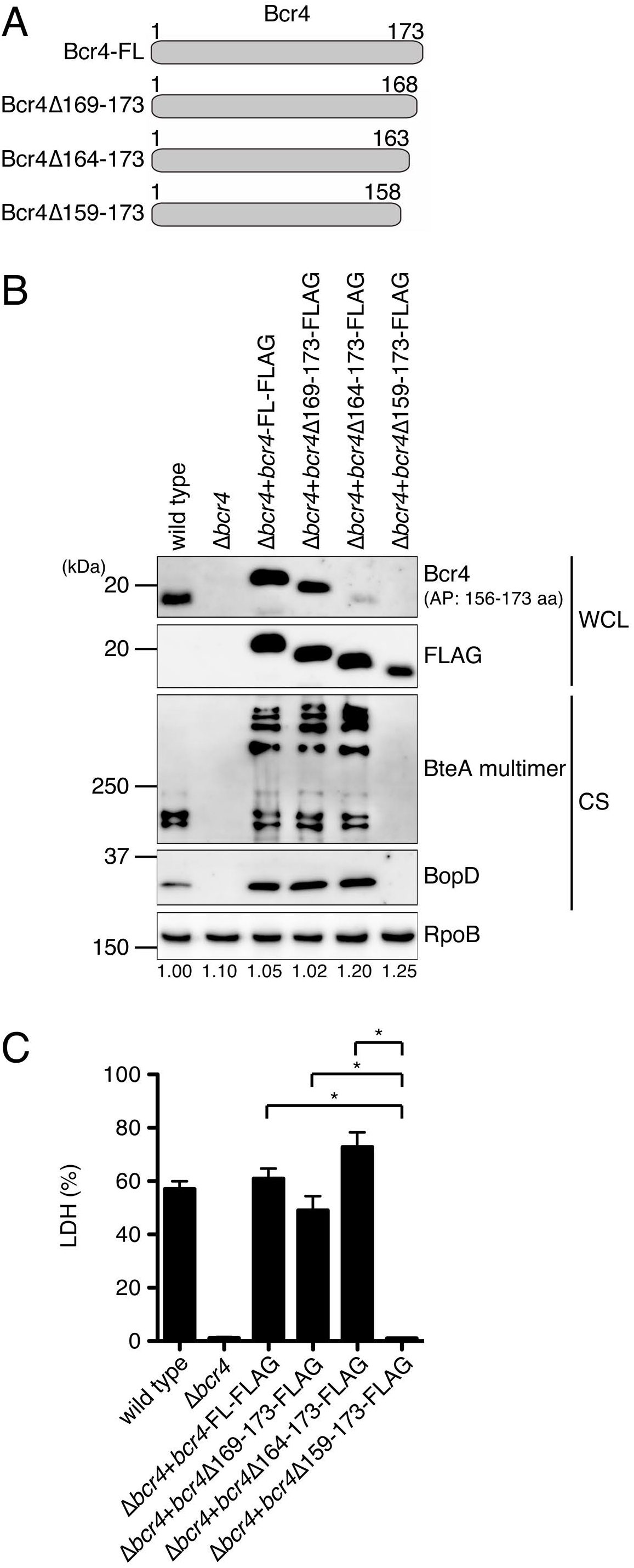
The Bcr4 domain required for T3SS activity. (A) Bcr4 derivatives used for the analysis of T3SS activity are depicted. (B) The *B. bronchiseptica* wild-type strain, Bcr4-deficient strain (Δ*bcr4*), and Δ*bcr4* producing Bcr4-FL-FLAG (Δ*bcr4*+*bcr4*-FL-FLAG), Bcr4Δ169-173-FLAG (Δ*bcr4*+*bcr4*Δ169-173-FLAG), Bcr4Δ164-173-FLAG (Δ*bcr4*+*bcr4* Δ164-173-FLAG), and Bcr4Δ159-173-FLAG (Δ*bcr4*+*bcr4*Δ159-173-FLAG) were cultured in SS medium. Whole cell lysates (WCL) and culture supernatant fraction (CS) samples were separated by SDS-PAGE and analyzed by Western blotting using antibodies against Bcr4, FLAG, BteA, BopD or RpoB. RpoB was used as an internal loading control. AP indicates the amino acids region of the antigen peptide used for anti-Bcr4 antibody generation. WCL and CS samples were prepared from equal volumes of bacterial culture. When we performed Western blotting using anti-BteA or BopD antibodies, we loaded a 100- or 50-fold smaller amount of Δ*bcr4*+*bcr4*-FL-FLAG, Δ*bcr4*+*bcr4*Δ169-173-FLAG and Δ*bcr4*+*bcr4*Δ164-173-FLAG CS samples on the SDS-PAGE gel than on the wild-type, Δ*bcr4* and Δ*bcr4*+*bcr4*Δ159-173-FLAG CS samples to avoid obtaining excess signal intensities. The numbers at the bottom of the lower panel indicate the relative signal intensity of RpoB measured using ImageJ software. (C) L2 cells were infected with each strain at an moi of 50 for 1 h. The amounts of LDH released into the extracellular medium from infected cells are shown, and the relative cytotoxicity (percent) was determined as described in the Materials and Methods. Error bars indicate the SEMs from triplicate experiments. \**P*<0.05. Experiments were performed at least three times, and representative data are shown.

### BscI is required for the function of the T3SS in *B. bronchiseptica*

BscI is a homologue of YscI, an inner rod (SctI) of the *Yersinia* T3SS, and YscI is essential for the function of the T3SS (22). To confirm that BscI is required for the function of the *B. bronchiseptica* T3SS, the *B. bronchiseptica* S798 wild-type, BscI-deficient strain (Δ*bscI*), BscI-complemented strain (Δ*bscI*+*bscI*), and BscN-deficient strain (Δ*bscN*) were cultured in SS medium. BscN is a protein predicted to function as an ATPase (SctN) and is required for the T3SS function in *Bordetella bronchiseptica* (Table S1) (6, 27). We used the Δ*bscN* as a T3SS-deficient strain. WCL and CS samples were prepared, and these samples were separated by SDS-PAGE and analyzed by Western blotting using antibodies against BscI, BteA, BopD, Bsp22 (SctA, a translocator and a component of the filament-like structure; Fig. 7) or FHA (filamentous hemagglutinin, an adhesion factor) (Fig. 4A). As expected, the FHA signals were detected in the WCL and CS samples of all strains used here (Fig. 4A). In contrast, no signals of BscI were detected in the WCL samples of all strains used here (Fig. 4A). The BscI signal was detected in the CS of Δ*bscI*+*bscI*, and a faint BscI signal was detected in the CS of the wild-type (Fig. 4A). In order to examine whether the signal we obtained around 20 kDa in the western blot using anti-BscI antibody in Fig. 4A was specific or nonspecific, we prepared the supernatant fraction from Δ*bsp22* strain. As a result, the signal disappeared in the supernatant fraction of the Δ*bsp22* strain (Fig. S3), suggesting that the signal around 20 kDa obtained in the western blot was a nonspecific interaction between anti-BscI antibody and an excess amount of Bsp22 on the membrane. The BteA, BopD and Bsp22 signals were detected in the CS of the wild-type and Δ*bscI*+*bscI*, but not in those of Δ*bscI* and Δ*bscN* (Fig. 4A). BspR is a negative regulator that represses the transcription of genes on the *bsc* locus (14). In *Bordetella* strains that are unable to secrete type III secreted proteins, such as Bcr4-deficient strains (Δ*bcr4*), the repression of the *bsc* locus transcriptions by BspR is enhanced (24). Therefore, we speculated that the production of BopD and Bsp22 was intensely suppressed by BspR in Δ*bscI* due to the loss of T3SS activity. To test whether this hypothesis is correct, we generated Δ*bspR*Δ*bscI* (a BspR- and BscI-deficient strain). WCL and CS samples were prepared from the *B. bronchiseptica* S798 wild-type strain, Δ*bspR*, Δ*bspR*Δ*bscI* or Δ*bspR*Δ*bscI*+*bscI* (BscI-complemented BspR- and BscI-deficient strain), and these samples were separated by SDS-PAGE and analyzed by Western blotting using antibodies against Bsp22 or BopB (SctE, a translocator and a type III secreted protein). As a result, the Bsp22 and BopB signals were detected in the WCL samples of the wild-type, Δ*bspR*, Δ*bspR*Δ*bscI* and Δ*bspR*Δ*bscI*+*bscI* strains (Fig. 4B). The Bsp22 and BopB signals were detected in the CS samples of the wild-type, Δ*bspR* and Δ*bspR*Δ*bscI*+*bscI* strains, but not in that of the Δ*bspR*Δ*bscI* strain (Fig. 4B). These results suggest that in *B. bronchiseptica*, BscI is required for the function of the T3SS and is secreted out of the bacterial cell.

**Fig. 4.**
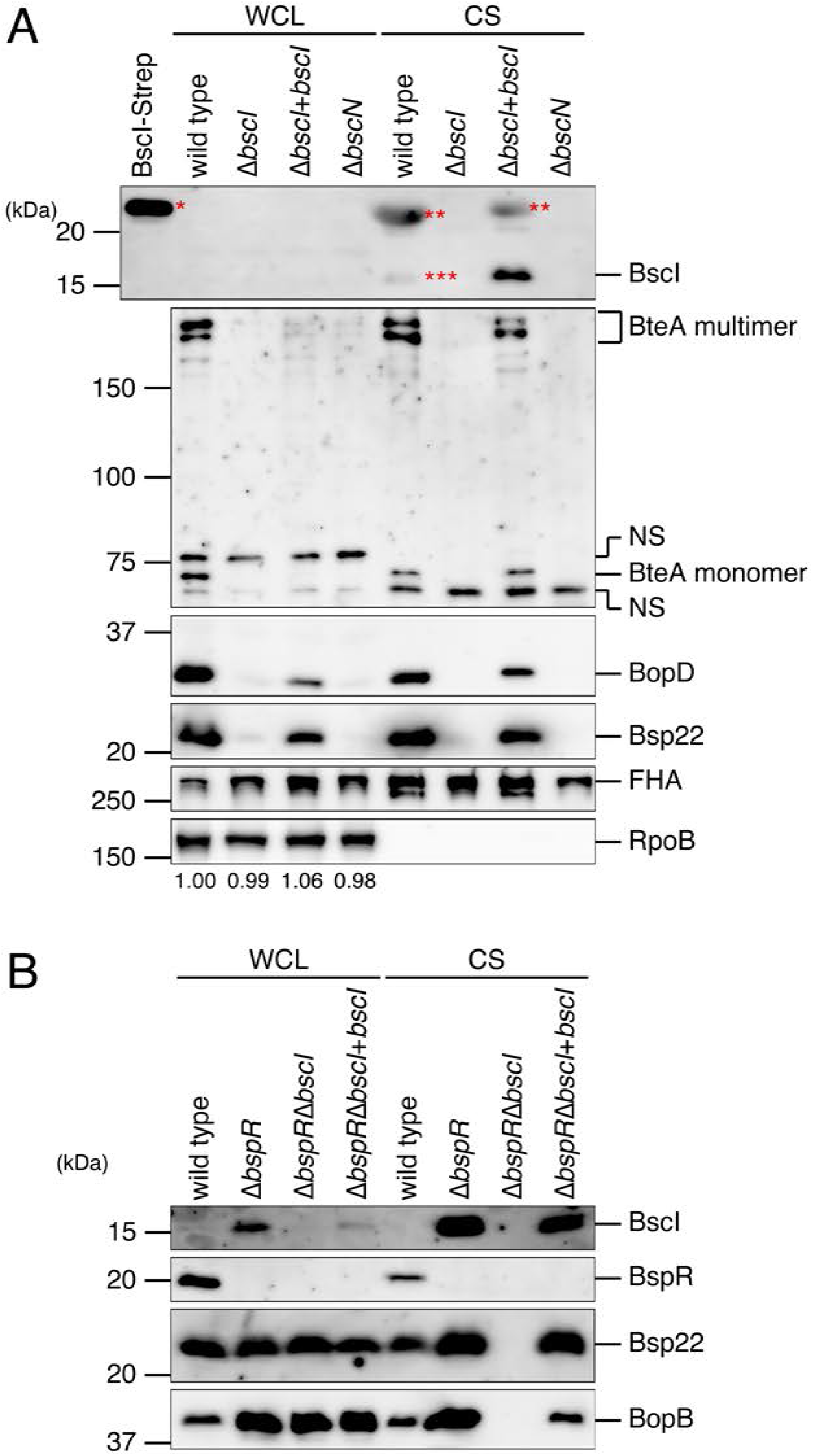
The effect of BscI deletion on T3SS activity in *B. bronchiseptica*. (A) Whole cell lysates (WCL) and culture supernatant fraction (CS) samples were prepared from the wild-type strain, Δ*bscI* (BscI-deficient strain), Δ*bscI*+*bscI* (BscI-complemented strain) and Δ*bscN* (T3SS-inactive strain) cultured in SS medium. WCL and CS samples were separated by SDS-PAGE and analyzed by Western blotting using antibodies against BscI, BteA, BopD, Bsp22, FHA or RpoB. BscI tagged with Strep (BscI-Strep) was loaded as a control. *BscI-Strep signal; **NS (nonspecific signals); ***A faint signal of BscI. The numbers at the bottom of the lower panel indicate the relative signal intensity of RpoB measured using ImageJ software. (B) WCL and CS were prepared from the wild-type, Δ*bspR* (BspR-deficient strain), Δ*bspR*Δ*bscI* (BspR- and BscI-deficient strain) and Δ*bspR*Δ*bscI*+*bscI* (BscI-complemented BspR- and Bcr4-deficient strain) cultured in SS medium. WCL and CS were separated by SDS-PAGE and analyzed by Western blotting using antibodies against BscI, BspR, Bsp22 or BopB antibodies. Loaded WCL and CS samples were prepared from equal volumes of bacterial culture. When we performed Western blotting using antibodies against Bsp22 or BopB, we loaded a 25- or 5-fold smaller amount of Δ*bspR*, Δ*bspR*Δ*bscI* and Δ*bspR*Δ*bscI*+*bscI* WCL samples, and 5- or 10-fold smaller amount of Δ*bspR* and Δ*bspR*Δ*bscI*+*bscI* CS samples on the SDS-PAGE gel than the wild-type to avoid obtaining excess signal intensities. Experiments were performed at least three times, and representative data are shown.

### BscI plays an important role in the function of the *B. bronchiseptica* T3SS during infection of cultured mammalian cells

To further investigate whether BscI is required for the construction and function of the T3SS in *B. bronchiseptica*, we infected L2 cells with the wild-type, Δ*bscI*, Δ*bscI*+*bscI* or Δ*bscN* and measured the number of Bsp22 (SctA, a translocator, and a component of the filament-like structure; Fig. 5) signals detected on L2 cells by immunofluorescence microscopy. For the efficient detection of Bsp22 signals, we cultured the bacteria in iron-depleted SS medium, which has been shown to increase the amount of type III secreted protein secreted by *B. bronchiseptica* (28). L2 cells were infected with bacteria for 1 h at an MOI of 2000. Then, Bsp22, F-actin and bacteria were stained with anti-Bsp22 antibody, Rhodamine-Phalloidin and DAPI, respectively (Fig. 5A). The DAPI fluorescence signals showing bacterial genomic DNA were detected to the same extent among the L2 cells infected with each bacterial strain, suggesting that the amounts of each mutant adhered to the cells were not significantly reduced when compared to the wild-type strain (Fig. 5A). The fluorescence signals of Bsp22 were detected on the L2 cells infected with the wild-type or Δ*bscI*+*bscI*, but not on the L2 cells infected with Δ*bscI* or Δ*bscN* (Fig. 5 A, B). We also infected L2 cells with the wild-type, Δ*bscI*, Δ*bscI*+*bscI* or Δ*bscN* for 1 h at an MOI of 200 and measured the amount of LDH released into the extracellular medium. The results showed that LDH was detected in the medium of L2 cells infected with the wild-type or Δ*bscI*+*bscI*, but not in the medium of L2 cells infected with Δ*bscI* or Δ*bscN* (Fig. 5 C). These results suggest that BscI is required for the construction of the T3SS and induction of T3SS-dependent death of mammalian cells in *B. bronchiseptica*.

**Fig. 5.**
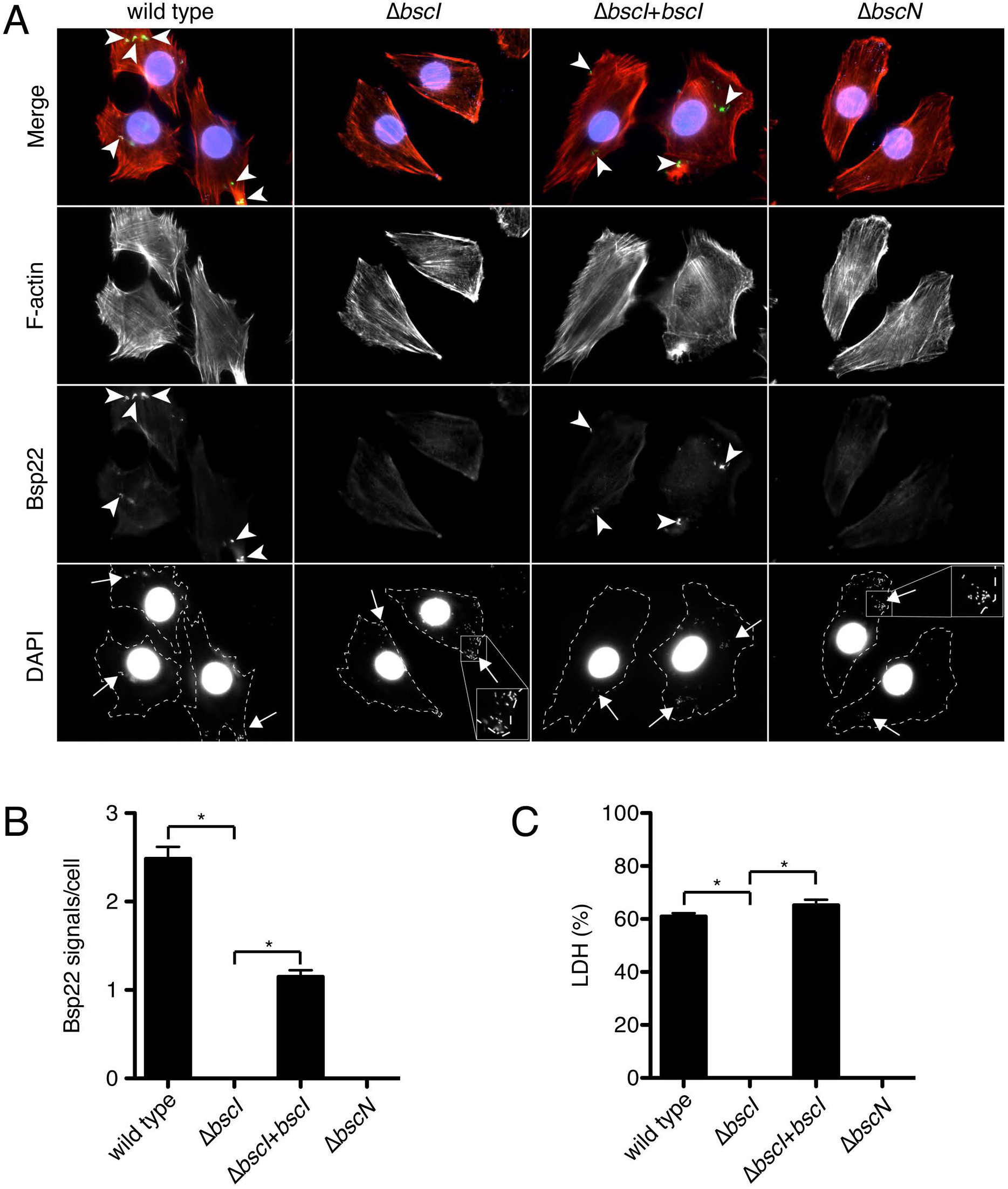
Immunofluorescent staining of Bsp22 on L2 cells infected with *B. bronchiseptica* and the results of the LDH assay. (A) L2 cells were infected with the wild-type strain, Δ*bscI* (BscI-deficient strain), Δ*bscI*+*bscI* (BscI-complemented strain) or Δ*bscN* (T3SS-inactive strain) as described in the Materials and Methods. After fixation, cells were stained with anti-Bsp22 antibody (green), rhodamine phalloidin (red) and DAPI (blue). The inset shows higher magnification of the boxed area in the image. Arrowheads and arrows indicate the signals of Bsp22 (green) and bacteria, respectively. (B) Bsp22 signals per one cell were counted under a fluorescent microscope. At least 120 cells were randomly chosen. (C) The results of LDH assay at an MOI of 200 are shown as a histogram. Error bars indicate the SEMs from triplicate experiments. \**P*<0.05. Experiments were performed at least three times, and representative data are shown.

### Bcr4 is required for the stability of BscI and is suggested to interact with BscI in *B. bronchiseptica*

As mentioned above, our results suggested that Bcr4 interacts with BscI (Fig. 1). Because Bcr4 has various properties like those of chaperones for the type III secreted proteins (25), e.g., low molecular mass (18.1 kDa) and low isoelectric point (pI 4.41), we investigated whether Bcr4 is involved in the stability of BscI. We cultured the *B. bronchiseptica* wild-type, Bcr4-deficient strain (Δ*bcr4*, T3SS-inactive strain) (24), BspR-deficient strain (Δ*bspR*, T3SS-overproducing and -oversecreting strain) (14) or BspR- and Bcr4-deficient strain (Δ*bspR*, a T3SS-overproducing but -inactive strain) (24) in SS medium. The prepared WCL samples were separated by SDS-PAGE, and Western blotting was performed using antibodies against BscI, BscJ (SctJ, a protein composing the inner membrane ring of the T3SS), or Bcr4. As shown in Fig. S1, the negative regulatory function of BspR, which represses the *bsc* locus transcription, is enhanced in the strain that loses the activity of T3SS, e.g., Δ*bcr4* (24). Because of this property of BspR, the BscJ signal of the WCL sample was weaker in Δ*bcr4* than in the wild-type (Fig. 6A). In Δ*bspR* and Δ*bspR*Δ*bcr4*, BscJ signals of the WCL samples were more strongly detected when compared to the wild-type (Fig. 6A) due to the BspR deficiency. On the other hand, the BscI signals of the WCL and CS samples were detected in Δ*bspR*, but not in the Δ*bspR*Δ*bcr4* strain (Fig. 6A). To investigate the presence of *bscI* mRNA in Δ*bspR*Δ*bcr4*, a quantitative RT-PCR was performed. The results showed that the amount of *bscI* mRNA in Δ*bspR*Δ*bcr4* was 1.5-fold higher than that of *bscI* in Δ*bspR* (Fig. S4), demonstrating that the *bscI* gene is transcribed in Δ*bspR*Δ*bcr4*. These results suggest that Bcr4 is necessary for the stability of BscI in *B. bronchiseptica*. Finally, we examined whether Bcr4 binds to BscI inside of the *B. bronchiseptica* cells. As described in the Discussion section, we were unable to detect an interaction of Bcr4 with BscI by immunoprecipitation because of the low level of BscI protein in bacterial cells. Therefore, we attempted to reveal the interaction by using a cross linker. We treated the wild-type, Δ*bspR* and Δ*bspR*Δ*bcr4* strains with disuccinimidyl suberate (DSS) and then prepared the whole cell lysate samples. The samples were separated by SDS-PAGE and then analyzed by Western blotting using anti-Bcr4 antibody. As a result, an extra signal around 33 kDa was evident in Δ*bspR* but not in Δ*bspR*Δ*bscI* (Fig. 6B). The size of the extra signal was almost equivalent to that of the Bcr4-BscI complex, suggesting the possibility that Bcr4 binds to BscI in the *B. bronchiseptica* cells.

**Fig. 6.**
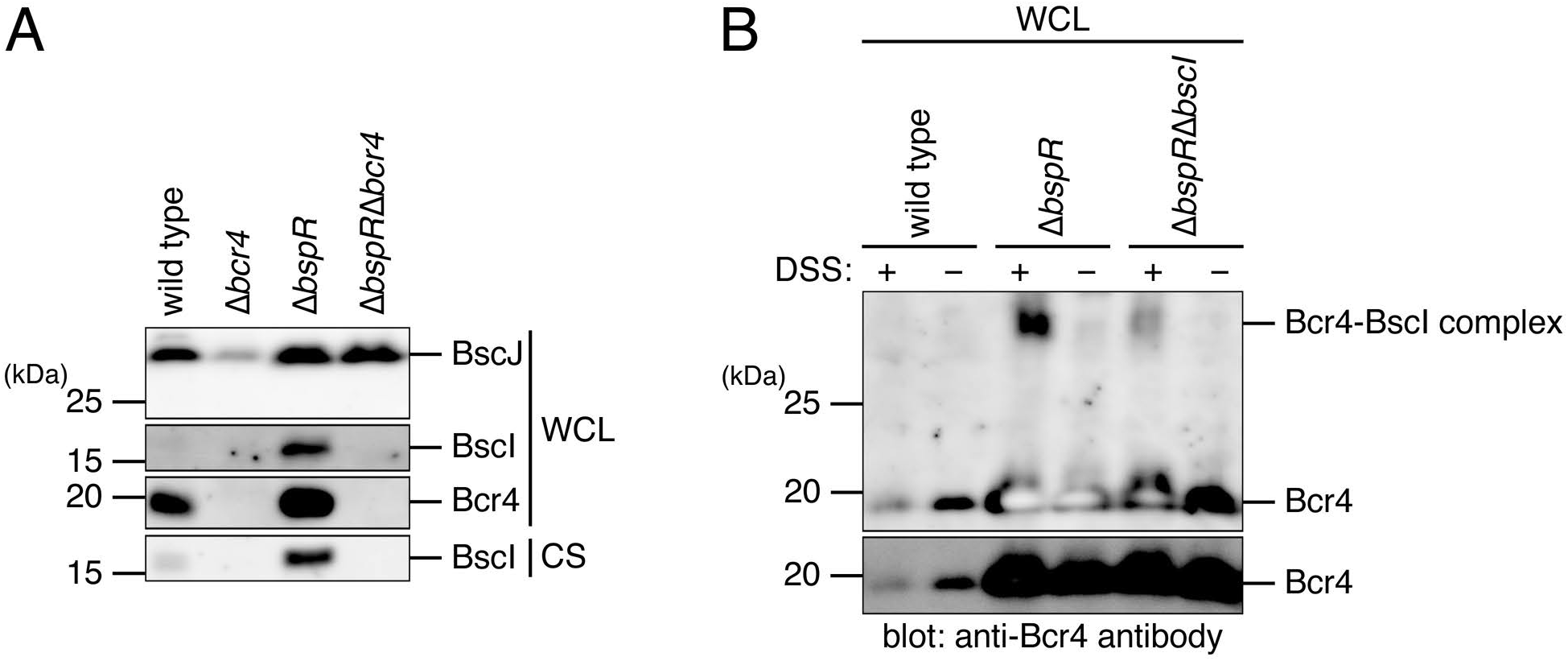
The effect of Bcr4 on BscI stability and the suggestive interaction of Bcr4 with BscI in *B. bronchiseptica*. (A) The whole cell lysates (WCL) and culture supernatant fraction (CS) were prepared from the wild-type strain, Δ*bcr4* (Bcr4-deficient strain), Δ*bspR* (BspR-deficient strain) or Δ*bspR*Δ*bcr4* (BspR- and Bcr4-deficient strain) cultured in SS medium. WCL and CS were separated by SDS-PAGE and analyzed by Western blotting with antibodies against BscI (inner-rod protein), BscJ (inner-ring protein) and Bcr4. The samples were prepared from equal volumes of bacterial culture. When we carried out Western blotting using anti-BscJ or Bcr4 antibodies, we loaded a 10-fold-smaller amount of Δ*bspR* or Δ*bspR*Δ*bcr4* WCL samples on the SDS-PAGE than WT or Δ*bcr4* WCL samples to avoid obtaining excess signal intensities. (B) WCL were prepared from the wild-type, Δ*bspR* or Δ*bspR*Δ*bcr4* treated with (+) or without (−) cross linker disuccinimidyl suberate (DSS). WCL were separated by SDS-PAGE and analyzed by Western blotting with anti-Bcr4 antibody. The lower panel is a short exposure of the upper panel. Experiments were performed at least three times, and representative data are shown.

## Discussion

In the present study, we found that deletion of BscI in *B. bronchiseptica* abolished the extracellular secretion of type III secreted proteins (Fig. 4, 5). In addition, when Bcr4 was deleted in the *B. bronchiseptica* BspR-deficient strain, Bscl (SctI, a protein composing the inner rod of the *Bordetella* T3SS) was not detected in the bacterial cells (Fig. 6A). These results suggest that Bcr4 contributes to the establishment of the T3SS in *B. bronchiseptica* by stabilizing BscI (Fig. 7).

**Fig. 7.**
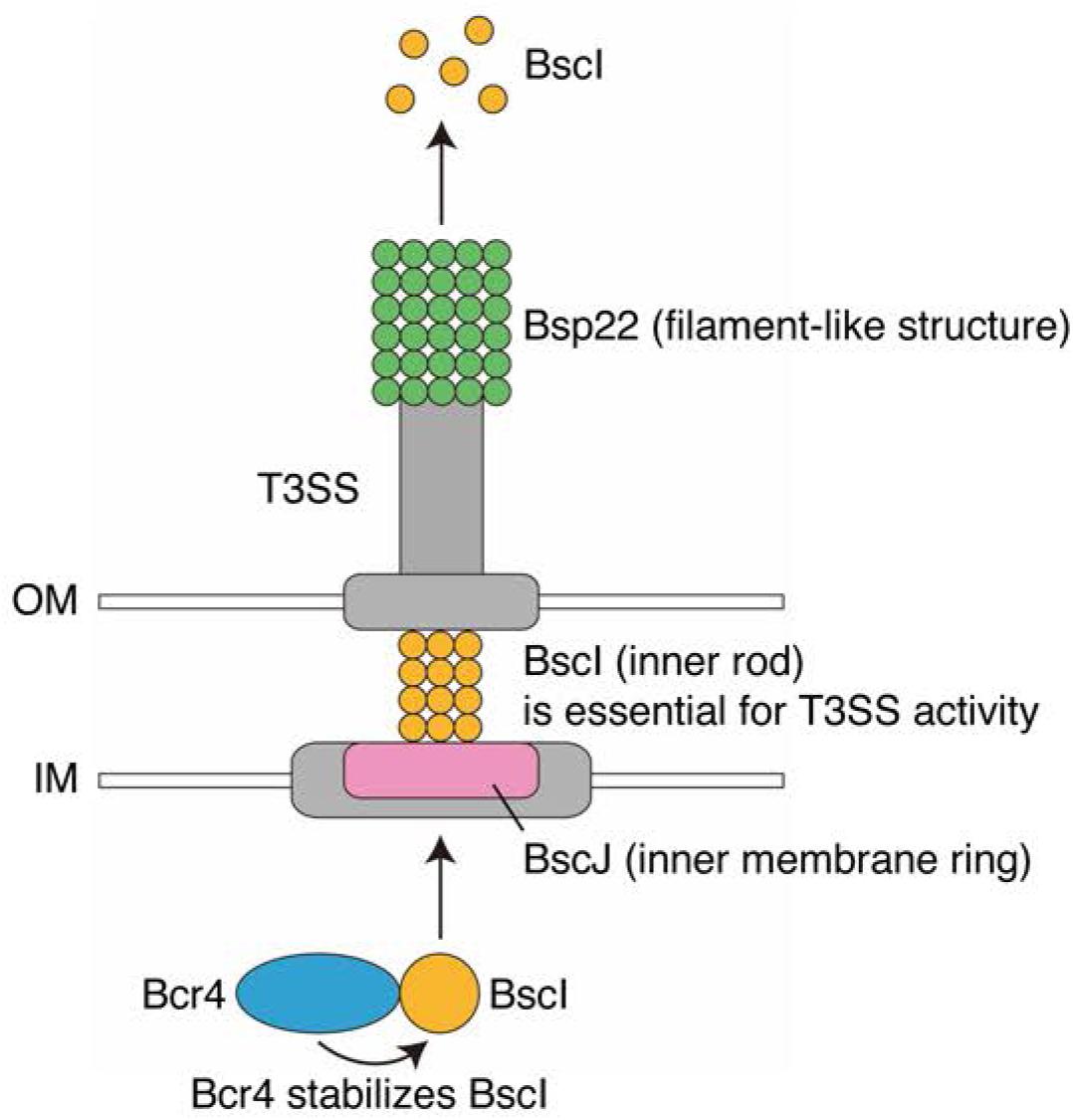
Schematic depiction of the Bcr4 and BscI functions. In the *B*. *bronchiseptica* bacterial cytosol, Bcr4 (blue) stabilizes BscI (orange). Then, BscI is presumed to be localized to the T3SS machinery (gray) and form the inner rod. BscI is essential for T3SS activity and secreted from the bacterial cell. BscJ (pink) is predicted to function as an inner membrane ring. Bsp22 (green) forms a filament-like structure. OM and IM indicate the outer membrane (white) and inner membrane (white), respectively.

To date, chaperones for needles, translocators and effectors have been reported in bacteria that retain the T3SS, such as *Yersinia* (29–32). However, a chaperone for the inner rod protein has not been reported, and the mechanism underlying stability of the inner rod in the bacterial cell and the transport of inner rods to the T3SS remain unclear.

In the present study, it was suggested that both the 58–109 and 110–173 amino acid regions of Bcr4 are required for the binding of Bcr4 to BscI (Fig. 2B). To investigate whether the 1–57 amino acids region of Bcr4 is also required for the binding of Bcr4 to BscI, we attempted to purify Bcr4 lacking the N-terminal 1–57 amino acids region (Bcr4Δ1-57-Strep). However, the amount of Bcr4Δ1-57-Strep produced in *E. coli* was extremely low (data not shown), so a pull-down assay could not be performed. Figure 3B suggests that the 159– 163 amino acids region of Bcr4 is required for the function of the T3SS. However, it is still unclear whether the 159–163 amino acids region of Bcr4 is required for binding to BscI. Further analysis is needed to clarify whether the binding of Bcr4 to BscI is required for the function of T3SS.

In this study, the BopD signal was detected in the WCL of the *B. bronchiseptica* wild-type strain, but the BscI signal was not detected (Fig. 4A). We detected *bscI* mRNA in *B. bronchiseptica*, and the amount of *bscI* mRNA was not less than that of *bopD* mRNA (Fig. S5). Therefore, the amount of BscI protein present in *B. bronchiseptica* was considered to be much lower than that of BopD. The reason for this finding is unknown, but it could be due to the low efficiency of translation of the *bscI* gene. In order to detect BscI in the whole cell lysate, we prepared the samples at 0 hr to 18 hr after suspending the bacteria in liquid broth. However, no signals were detected in the whole cell lysate samples prepared from the wild-type strain or the Δ*bcr4* strain (Fig. S6). We also used the Δ*bspR* and Δ*bspR*Δ*bcr4* strains, and detected BscI signals in the Δ*bspR* strain, but not in the Δ*bspR*Δ*bcr4* double knockout strain (Fig. S6), strongly suggesting that Bcr4 is important for the stability of BscI. In this study, we attempted to detect the BscI signal using a BspR-deficient strain (Δ*bspR*), in which the gene transcriptions on the *bsc* locus are promoted (Fig. 6), because the BscI signal was not detected in the WCL sample of the *B. bronchiseptica* wild-type (Fig. 4). The *bscJ* gene is located downstream of the *bscI* gene (Fig. 1A), and the BscJ protein is predicted to function as an inner membrane ring (SctJ), a component of the T3SS. The signal intensity of BscJ was not decreased in the WCL sample of the BspR/Bcr4 double-deficient strain (Δ*bspR*Δ*bcr4*) compared to that of Δ*bspR* (Fig. 6A), suggesting that Bcr4 is not involved in the stability of BscJ. Therefore, it is suggested that Bcr4 specifically stabilizes BscI. In order to detect the interaction between Bcr4 and BscI in the *B. bronchiseptica* cells, immunoprecipitation was performed using anti-BscI antibody against Δ*bspR* lysate. The immunoprecipitated fractions were analyzed by Western blotting using anti-BscI antibody; however, an evident BscI signal was not detected (data not shown). The results of our immunoprecipitation assay were thus unable to demonstrate a specific interaction between Bcr4 and BscI. We speculate that *B. bronchiseptica* did not maintain a sufficient amount of BscI in whole cell lysate to detect the interaction by immunoprecipitation. In Fig. 6B, we obtained an extra signal around 33 kDa in Western blotting using anti-Bcr4 antibody when we analyzed the lysate of *B. bronchiseptica* treated with a cross-linker, DSS. When we performed the Western blotting using anti-BscI antibody for the cross-linked lysate, the extra signal was not detected (data not shown).

In order to examine whether Bcr4 has structural similarity to any chaperones for the type III secreted proteins produced by other bacteria, we used AlphaFold2. As a result, we detected significant structural similarities to the other chaperones, *Aeromonas* AcrH (34) and *Pseudomonas* PscG (35) (Fig. S7). Although we obtained no plausible structural model for the interaction between BscI and Bcr4, the results strongly suggest that Bcr4 is a chaperone.

It is still unclear how Bcr4 antagonizes the BspR-negative regulation activity, and how overexpression of Bcr4 promotes the production and secretion of type III secreted proteins in *B. bronchiseptica*. Further analysis is needed to elucidate the detailed molecular mechanism by which Bcr4 contributes to the regulation and establishment of the T3SS.

## Materials and Methods

### Bacterial strains and cell culture

The strains used in this study are listed in Table 1. *B. bronchiceptica* S798 (6) was used as the wild-type strain. Δ*bscN* (a BscN-deficient strain), Δ*bcr4* (a Bcr4-deficient strain), Δ*bspR* (a BspR-deficient strain) and Δ*bspR*Δ*bcr4* (a BspR and Bcr4-deficient strain) have been described elsewhere (6, 14, 24). *Escherichia coli* DH10B, BL21 (36) and Sm10λ*pir* were used for DNA cloning, recombinant protein expression and conjugation, respectively (Table 1). *B. bronchiceptica* was grown on Bordet-Gengou agar plates at 37°C for 2 days. Fresh colonies of *B. bronchiceptica* were cultured in Stainer-Scholt (SS) medium (37) with a starting A_600_ of 0.23 under vigorous shaking conditions at 37°C. L2 cells, a rat lung epithelial cell line (ATCC CCL-149), were maintained in F-12K (Invitrogen). The cell culture medium was supplemented with 10% fetal calf serum (FCS). L2 cells were grown at 37°C under a 5% CO_2_ atmosphere.

**Table 1.**
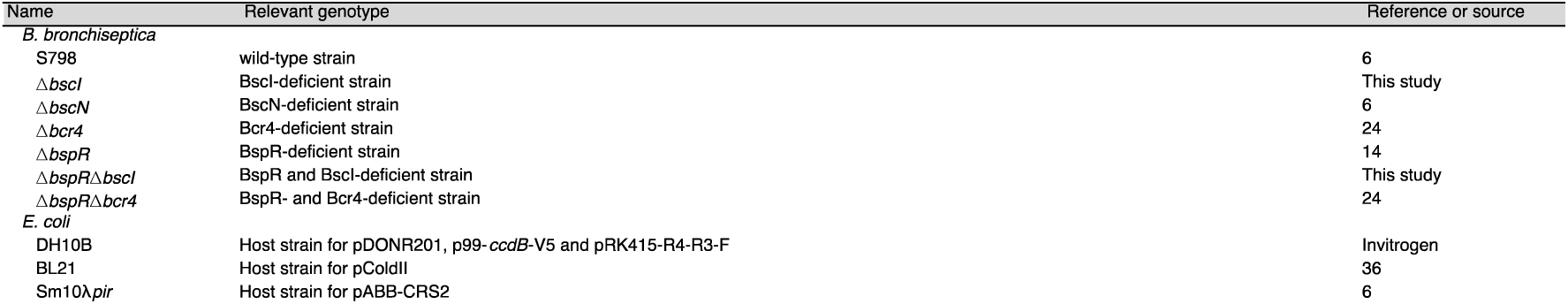
Bacterial strains used in the study

### Preparation of proteins from culture supernatants and whole bacterial cell lysates

Proteins secreted into bacterial culture supernatants and whole bacterial cell lysates were prepared as described previously (38). The loaded sample amounts were adjusted by the A_600_ of each bacterial culture to load samples prepared from the same amounts of bacteria. The protein samples were separated by SDS-PAGE and analyzed by Western blotting.

### Antibodies

Anti-BteA, anti-BopD, anti-Bsp22, BspR, BopB, and anti-Bcr4 antibodies were purified from rabbit serum in our previous study (6, 7, 10, 14, 24, 39). Mouse anti-V5 and anti-RNA polymerase beta subunit (RpoB) monoclonal antibodies were purchased from Santa Cruz Biotechnology and BioLegend, respectively. To detect filamentous hemagglutinin (FHA) signals, we used mouse anti-FHA serum kindly gifted from K. Kamachi (National Institute of Infectious Diseases). To prepare the anti-BscI and anti-BscJ antibodies, the peptides corresponding to the C-terminus regions of BscI (CKAIGRATQNVDTLARMS) and BscJ (CRGEGRGGAGAGATEGAGHD) were conjugated with hemocyanin from keyhole limpets (Sigma), respectively, by using 3-maleimidobenzoic acid N-hydroxysuccinimide ester (Sigma). These cross-linked peptides were used to immunize rabbits, and the resulting anti-sera were incubated with the same peptides immobilized on epoxy-activated sepharose 6B (Amersham) to obtain specific Ig-fractions.

### Immunofluorescence staining of L2 cells infected with *B. bronchiseptica*

Immunofluorescent staining assay was performed as previously described (38), with slight modifications. Briefly, L2 cells were seeded on coverslips in 6-well plates and incubated overnight. In order to detect adequate amounts of Bsp22 signals, we infected the L2 cells with *B. bronchiseptica* cultured in iron-depleted SS medium at an MOI of 2000. To avoid excessive killing of L2 cells with bacteria, the plates were not centrifuged. After being incubated for 1 h at 37°C under a 5% CO_2_ atmosphere, the infected L2 cells were immunostained. Average numbers of Bsp22 signals on single L2 cells were measured by a fluorescence microscope.

### LDH Assays

LDH assays were performed as previously described (38). Briefly, to examine whether lactate dehydrogenase (LDH) is released from *B. bronchiseptica*-infected cells, 5.0×10^4^ cells/well of L2 cells seeded in 24-well plates were infected with bacteria at an MOI of 50 or 200. The plates were centrifuged at 900×*g* for 5 min and incubated for 3 h at 37°C under a 5% CO_2_ atmosphere. The amounts of LDH were measured spectrophotometrically using a Cyto-Tox 96 Non-radioactive Cytotoxicity Assay kit (Promega). The LDH (%) level is shown as a ratio, with the value obtained from the well treated with 0.1% Triton X-100 set as 100%.

### Cross-linking Assay

The cross-linking assay was performed as previously described (40). A 100 µL culture of *B. bronchiseptica* was centrifuged at 20,000×*g* and the supernatant was removed. The pellet was washed with PBS and resuspended in 100 µL of PBS. Then, disuccinimidyl suberate (DSS (Thermo)) dissolved in DMSO was added at a final concentration of 10 mM. After incubation on ice for 1 h, Tris-HCl pH8.0 was added at a final concentration of 50 mM. The solution was centrifuged at 20,000×*g*, and the supernatant was removed. The pellet was dissolved in 2×SDS-PAGE sample buffer. The samples were prepared from the same amounts of bacteria based on the A_600_ values of the bacterial culture, then separated by SDS-PAGE, and analyzed by Western blotting.

### Statistical analyses

The statistical analyses were performed using the nonparametric test with a two-tailed p*-*value with Prism ver. 5.0f software (Graphpad, La Jolla, CA). Values of p<0.05 were considered significant.

### Others

The pull-down assay, the construction of *bscI* gene-disrupted or *bspR*/*bscI* double strains, the construction of the plasmids used for producing Bcr4 derivatives and BscI complementation, the quantitative reverse transcription-PCR, and Protein structure prediction are described in Text S. Plasmids and Primers used in this study are listed in Table S2 and S3, respectively.

## Acknowledgements

This work was supported in part by grants (19K07542 to A.A.; 20K07485 to A.K.; 19K07561 to T.H.) from the Ministry of Education, Culture, Sports, Science and Technology and the Japan Society for the Promotion of Science (KAKENHI), and a grant (JP22gm1610003 to M.S. and A.K.) from the Japan Agency for Medical Research and Development (AMED).

The funders had no role in the study design, date collection or analysis, the decision to publish, or the preparation of manuscript.

## Supplemental material

### Text S. Supplementary Information

**Fig. S1.**
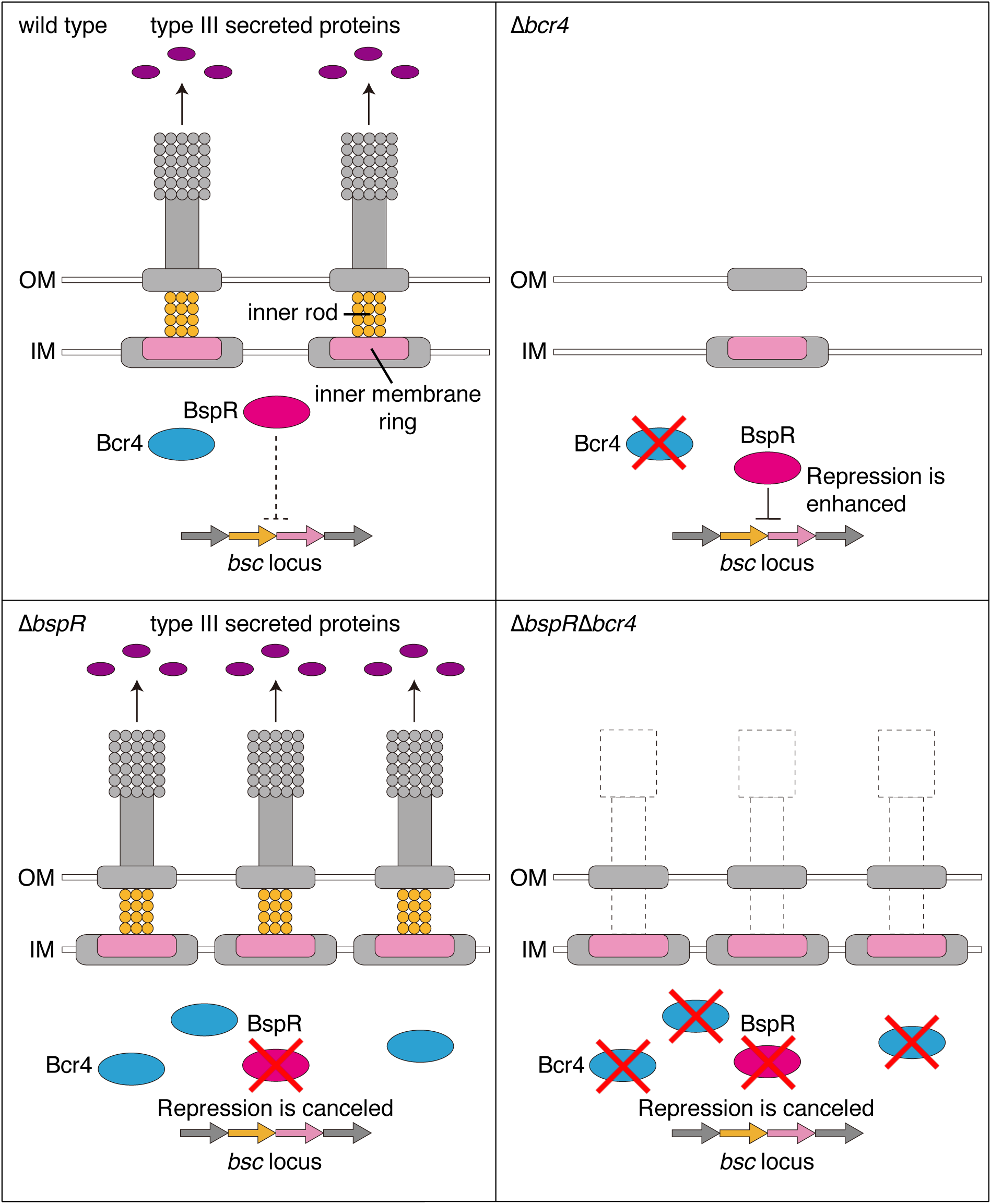
Construction of the T3SS machinery in strains lacking Bcr4 and/or BspR. In the *B. bronchiseptica* wild-type (upper left), the BspR negative regulation level for the *bsc* locus transcription is moderate, and the T3SS machinery is established. In the Bcr4-deficient strain (upper right), BspR strongly represses the *bsc* locus transcription, and construction of the T3SS machinery is incomplete. In the BspR-deficient strain (lower left), the negative regulatory effect of BspR is cancelled, and the construction of the T3SS machinery is promoted. In the BspR/Bcr4 double-deficient strain (lower right), while the *bsc* locus transcription is promoted because of BspR deficiency, T3SS is not functional.

**Fig. S2.**
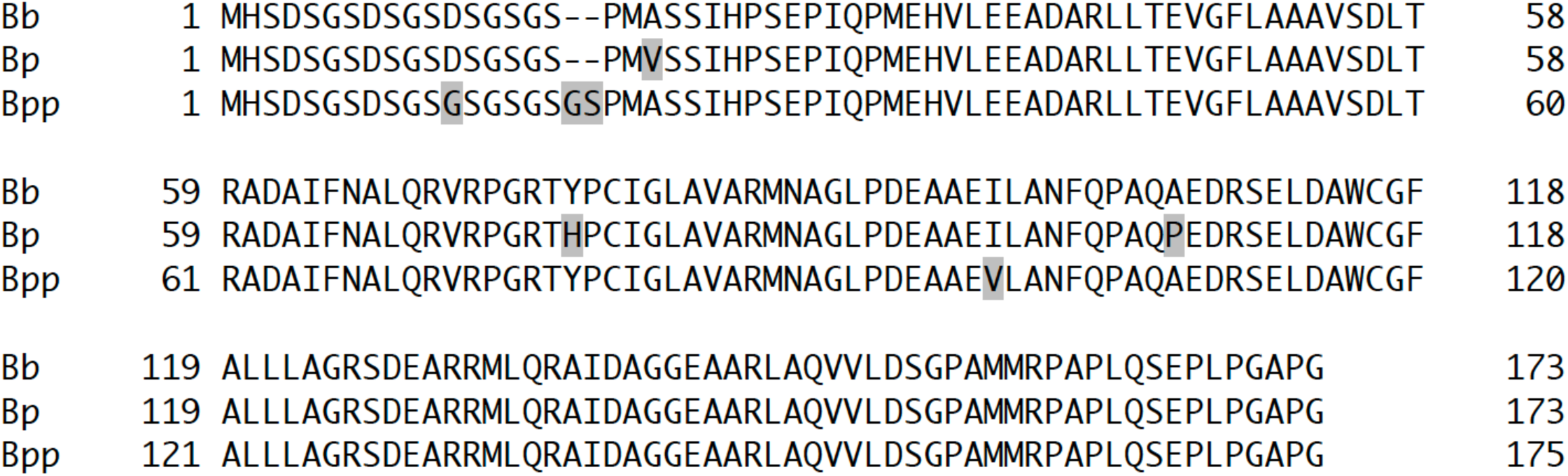
Alignment of Bcr4 amino acid sequences in representative *Bordetella* species. Bcr4 amino acid sequences of *B. bronchiseptica* S798 (Bb), *B. pertussis* Tohama I (Bp) and *B. parapertussis* 12822 (Bp) were compared using ClustalW. The grey highlighted letters represent amino acid residues that were different from those in Bb. Bcr4 of Bp and Bpp have 98.3% and 97.1% identities with those of Bp, respectively.

**Fig. S3.**
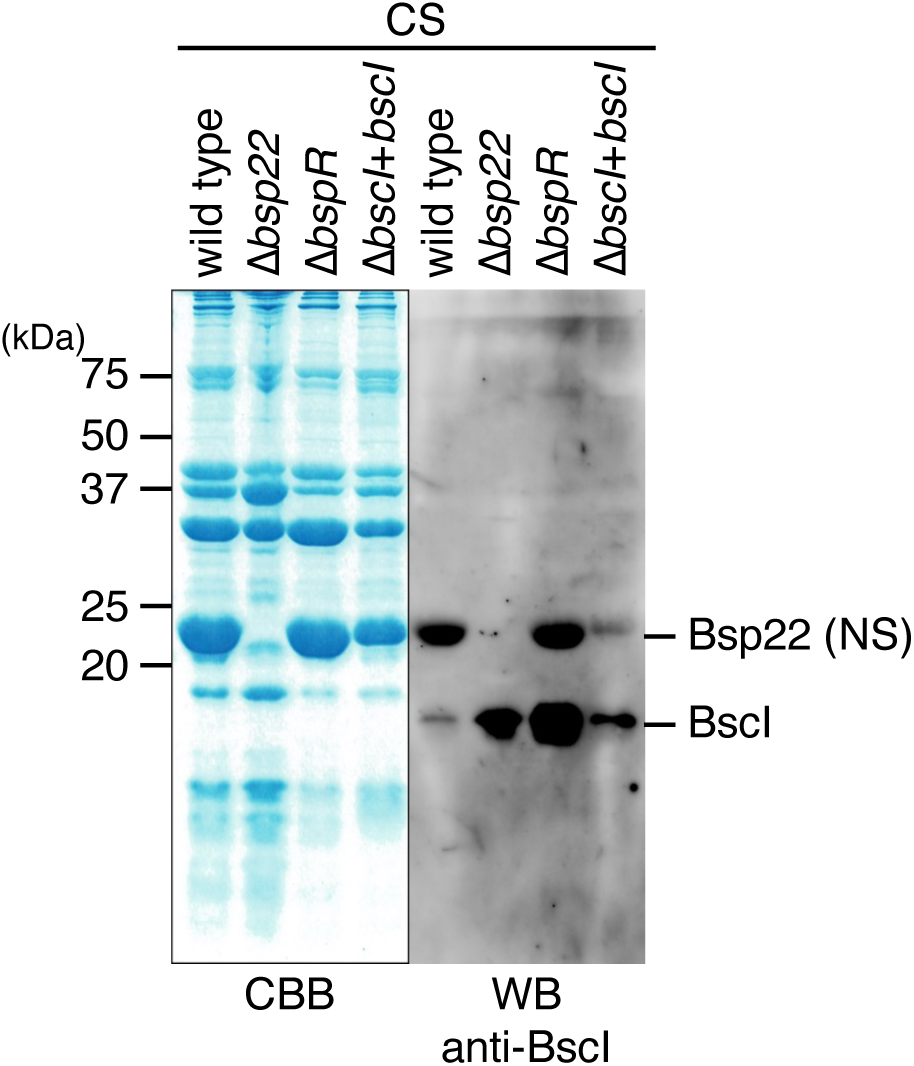
The nonspecific reaction of anti-BscI antibody to Bsp22. The culture supernatants (CS) were prepared from the wild-type strain, Δ*bsp22* (Bsp22-deficient strain), Δ*bspR* (BspR-deficient strain) or Δ*bscI*+*bscI* (BscI-complemented strain) cultured in SS medium. The CS samples were separated by SDS-PAGE and stained with Coomassie Brilliant Blue (CBB, left panel) or analyzed by Western blotting (WB) with anti-BscI antibody (right panel). NS indicates nonspecific signals. Experiments were performed at least three times, and representative data are shown.

**Fig. S4.**
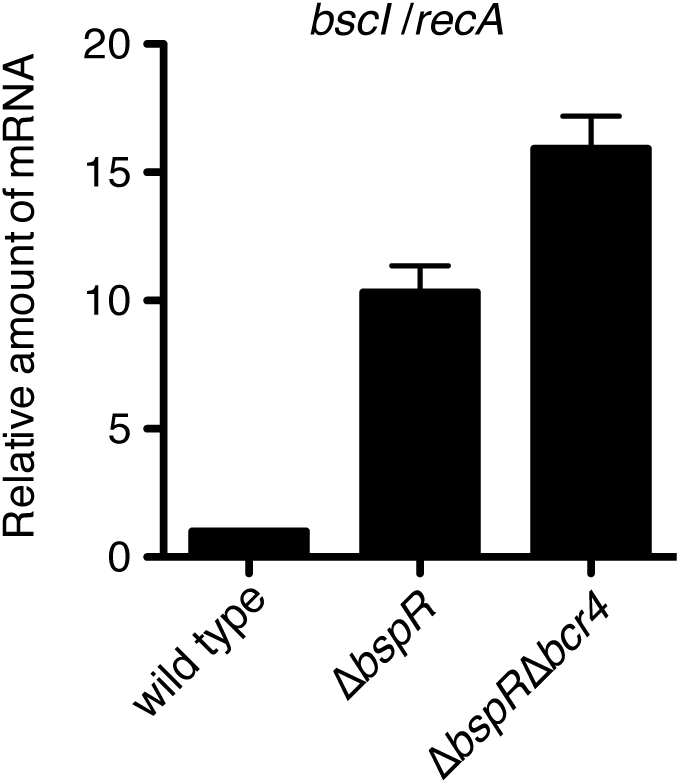
The results of the RT-PCR analysis for the mRNA level of *bscI* in *B. bronchiseptica* strains. Total RNA was prepared from the wild-type strain, Δ*bspR* (BspR-deficient strain) or Δ*bspR*Δ*bcr4* (BspR- and Bcr4-deficient strain) cultured in SS medium and subjected to a quantitative RT-PCR analysis. The histogram shows the relative amount of *bscI* mRNA normalized by the housekeeping gene, *recA* mRNA. Experiments were performed at least three times, and representative data are shown.

**Fig. S5.**
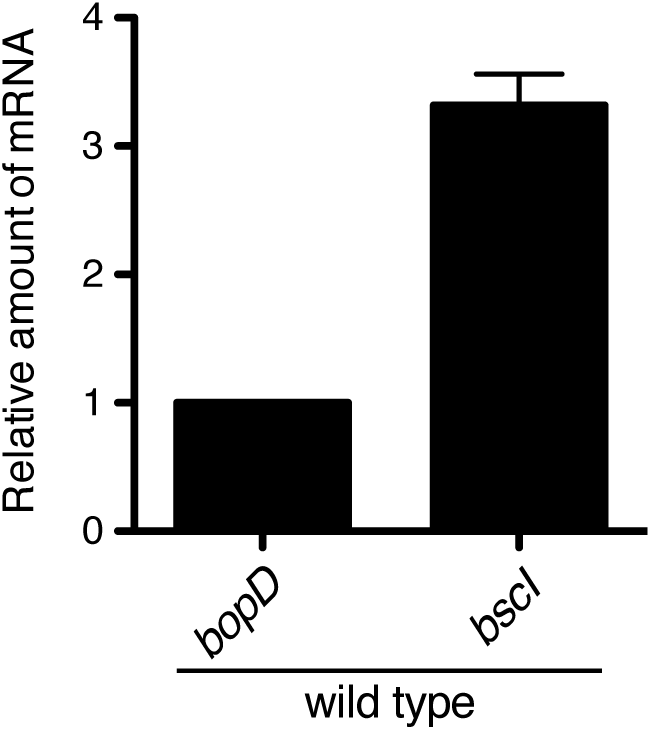
The results of the RT-PCR analysis for mRNA levels of *bopD* and *bscI* in the wild-type *B. bronchiseptica*. Total RNA was prepared from the wild-type strain cultured in SS medium and subjected to a quantitative RT-PCR analysis. The histogram shows the relative amount of *bopD* and *bscI* mRNA in the wild-type. The relative ratio of *bscI* mRNA is shown when the *bopD* mRNA amount is set as 1. Experiments were performed at least three times, and representative data are shown.

**Fig. S6.**
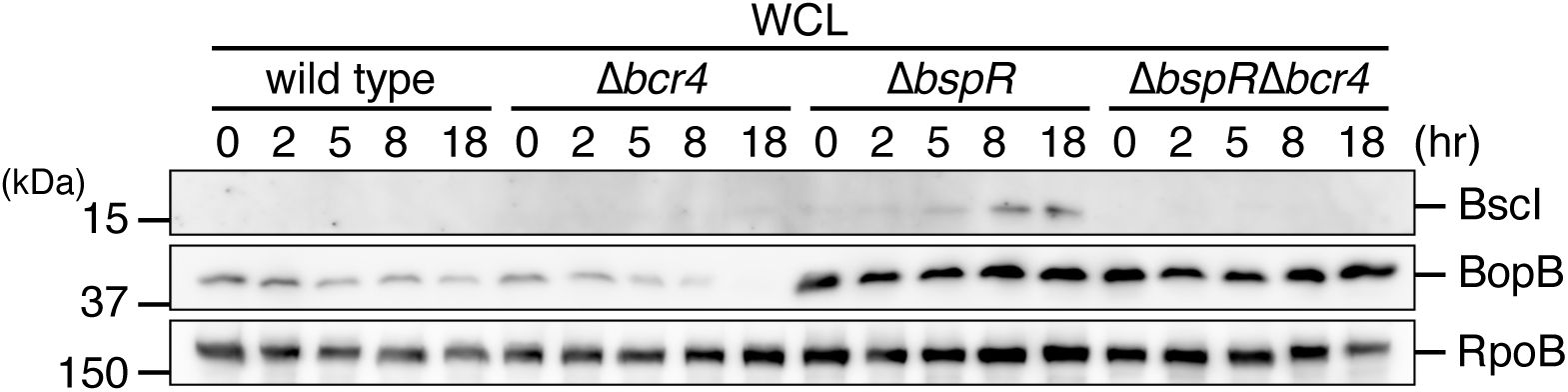
The time course of BscI production in *B. bronchiseptica*. The whole cell lysates (WCL) were prepared from the wild-type strain, Δ*bcr4* (Bcr4-deficient strain), Δ*bspR* (BspR-deficient strain) or Δ*bspR*Δ*bcr4* (BspR- and Bcr4-deficient strain) cultured in SS medium for 0, 2, 5, 8 or 18 hr. The WCL were separated by SDS-PAGE and analyzed by Western blotting with antibodies against BscI, BopB and RpoB. Experiments were performed at least three times, and representative data are shown.

**Fig. S7.**
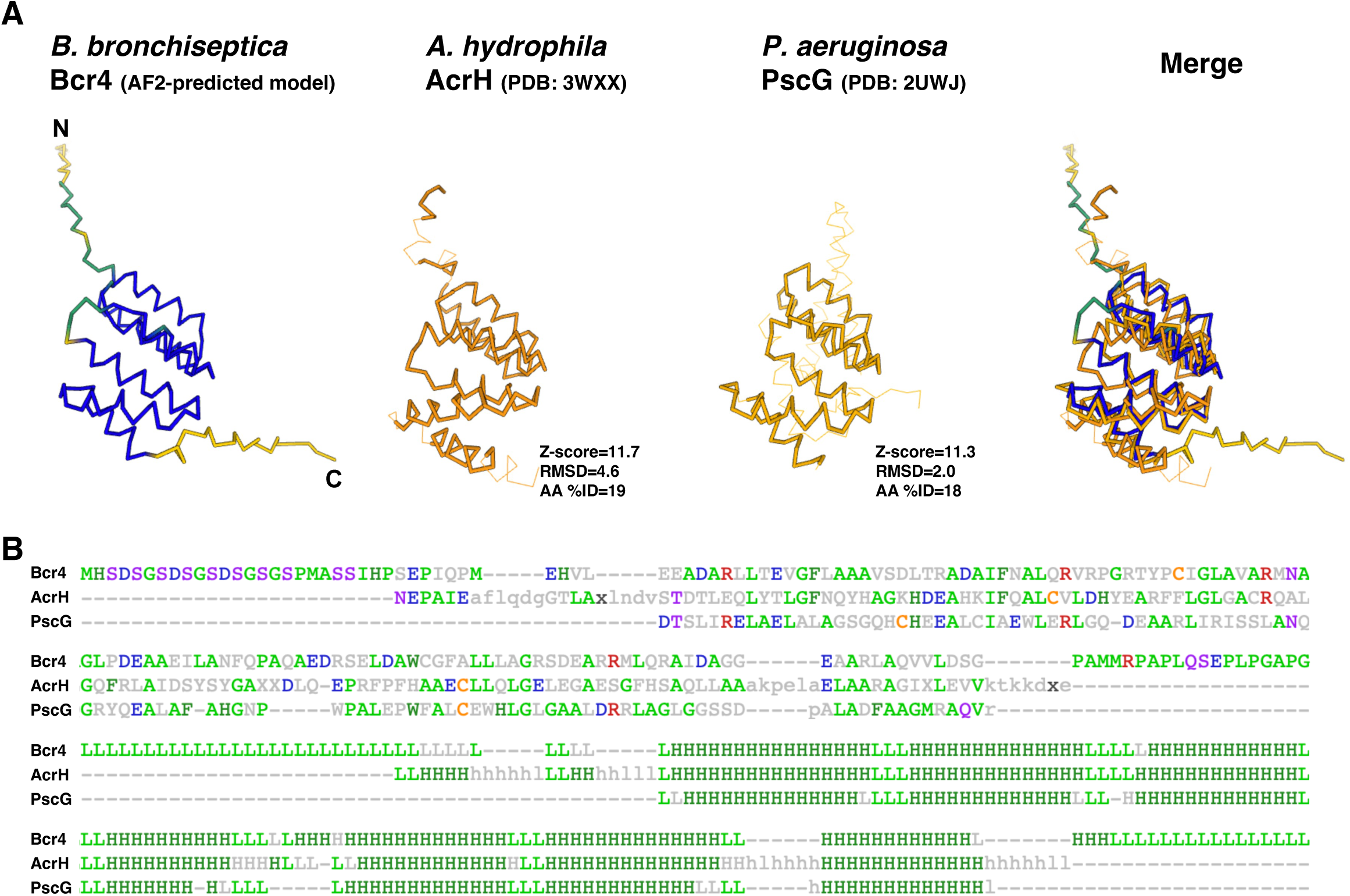
The predicted structural model of *B. bronchiseptica* Bcr4. (A) The AlphaFold2 (AF2)-predicted structural model of *B. bronchiseptica* Bcr4 and structural comparison with *Aeromonas hydrophila* AcrH (PDB: 3WXX) and *Pseudomonas aeruginosa* PscG (PDB: 2UWJ). Z-score, root mean square deviation (RMSD), and amino acid identity (AA %ID) of AcrH and PscG compared with Bcr4 are shown, respectively. (B) The pairwise sequence alignment of Bcr4, AcrH, and PscG. The most frequent amino acid type is colored. The secondary structure assignments (H/h: helix, E/e: strand, L/l: coil) are also shown.

**Table S1.**
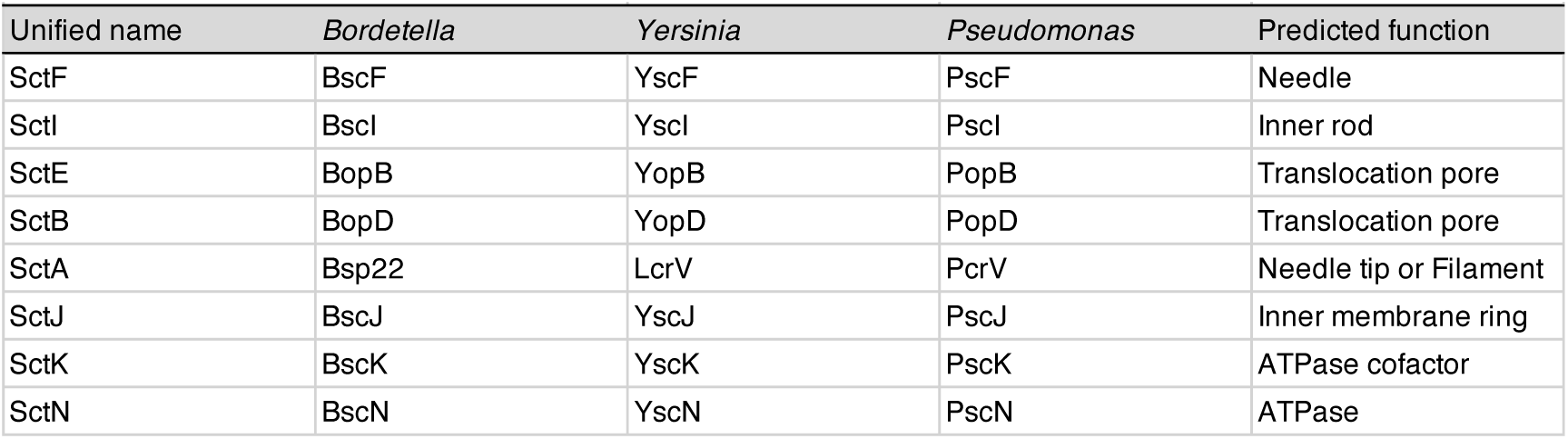
Nomencalture of the *Bordetella* T3SS component.

**Table S2.**
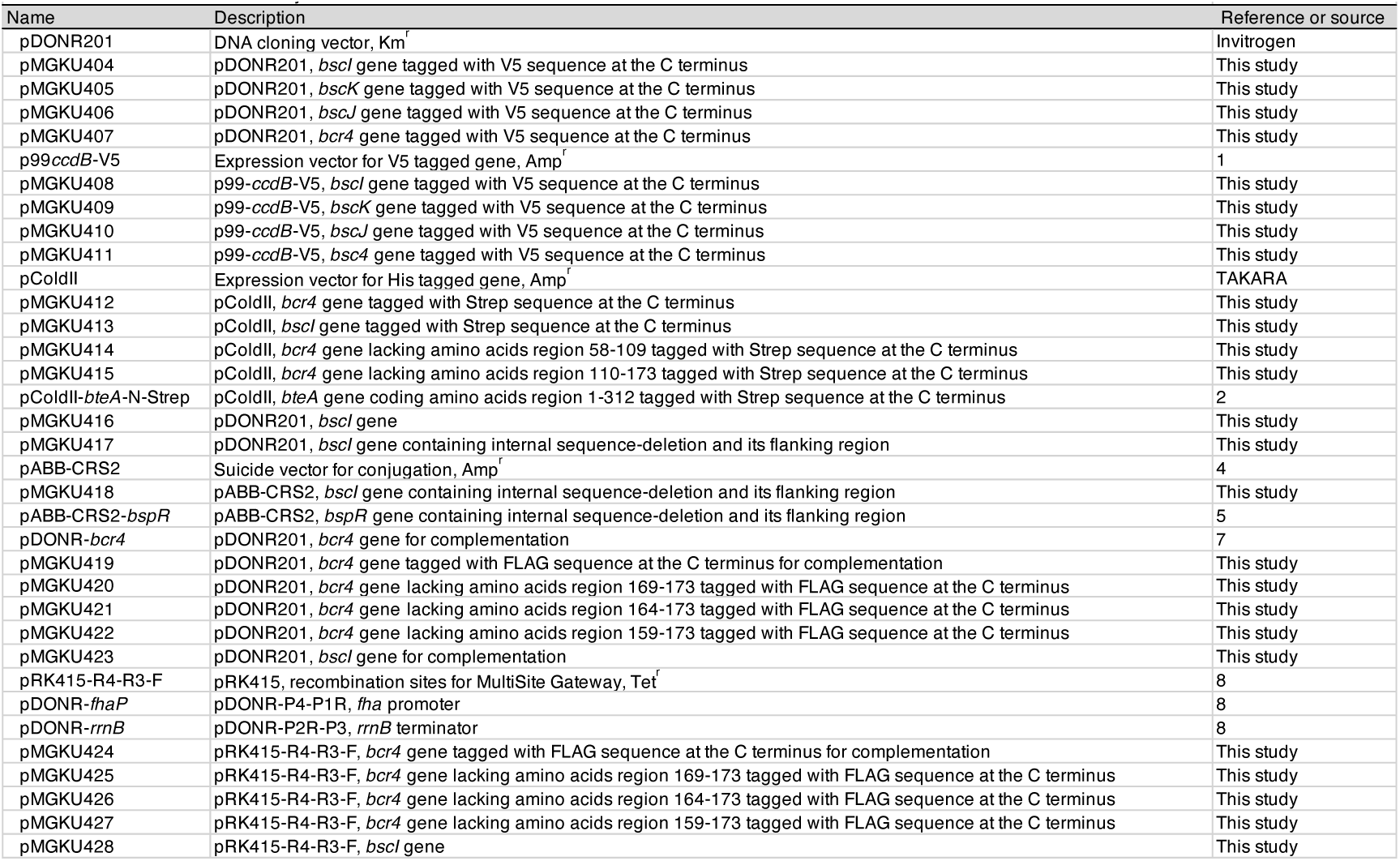
Plasmids used in the study.

**Table S3.**
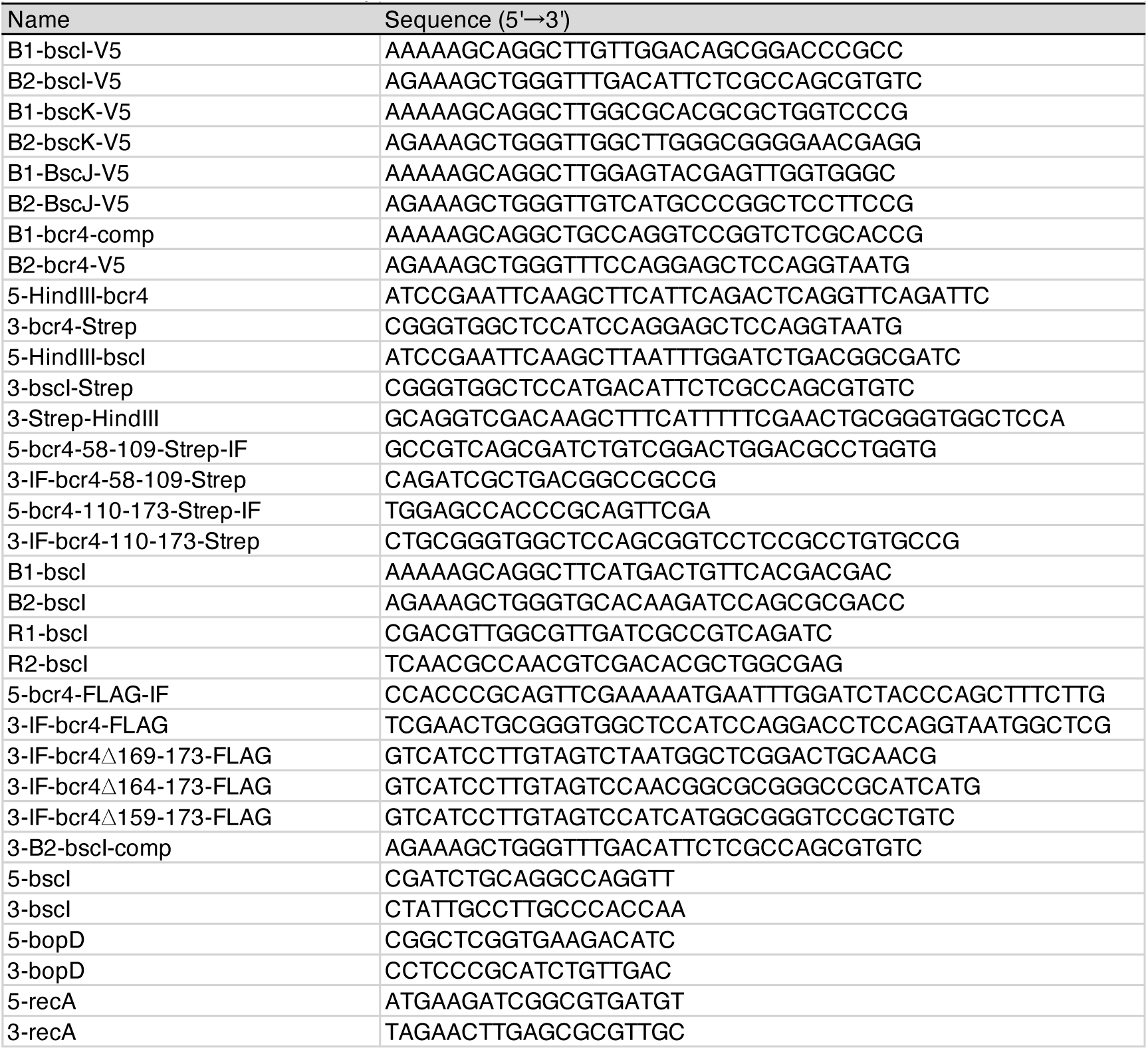
Primers used in the study.

## Supplementary Information

### SI Materials and Methods

#### Pull-down assay

The plasmids and primers used in this study are listed in Table S1 and S2, respectively. In order to express *bscI*, *bscK*, *bscJ* or *bcr4* tagged with a V5 sequence at the respective C-terminus (*bscI*-V5, *bscK*-V5, *bscJ*-V5 or *bcr4*-V5), we amplified DNA fragments encoding the *bscI*, *bscK*, *bscJ* or *bcr4* genes with the primer sets of B1-bscI-V5 and B2-bscI-V5, B1-bscK-V5 and B2-bscK-V5, B1-bscJ-V5 and B2-bscJ-V5 or B1-bcr4-comp and B2-bcr4-V5, respectively, using *B. bronchiseptica* S798 genomic DNA as the template. Each resulting PCR product was cloned into pDONR201 to obtain pMGKU404, pMGKU405, pMGKU406 or pMGKU407, respectively, by means of adapter PCR and site-specific recombination techniques using the Gateway cloning system (Invitrogen). Each plasmid was mixed with an expression vector such as p99*ccdB*-V5 (1) to obtain pMGKU408, pMGKU409, pMGKU410 or pMGKU411 using the Gateway cloning system.

BL21 cells carrying pMGKU408, pMGKU409 pMGKU410 or pMGKU411 were cultured overnight at 37°C with shaking, and then diluted 1:40 in LB liquid medium containing 50 µL/mL ampicillin and incubated for 2 h at 37°C with shaking. Each bacterial culture was further incubated for 5 h at 30°C in the presence of isopropyl-beta-thiogalactopyranoside (IPTG) at the final concentration of 1 mM. Bacteria were collected by centrifugation at 2,600×*g* for 15 min, and suspended in cold TBS containing protease inhibitor cocktail, cOmplete (Roche). Each bacterial suspension was sonicated, and each supernatant was used for the pull-down assay.

Next, in order to purify Bcr4 or BscI tagged with six histidine residues (6×His) at the respective N-terminus and Strep at the respective C-terminus, we amplified DNA fragments encoding *bcr4* or *bscI* with the primer sets of 5-HindIII-bcr4 and 3-bcr4-Strep, or 5-HindIII-bscI and 3-bscI-Strep using *B. bronchiseptica* S798 genomic DNA as the template. Each amplified DNA fragment was used as a template for 2nd PCR with a primer set consisting of the upper primer used in the 1st PCR and 3-Strep-HindIII to add a 24 bp sequence encoding the Strep tag. Each resulting PCR product was cloned into the HindIII recognition sites of pColdII to obtain pMGKU412 or pMGKU413 by the In-Fusion Cloning System (Clontech), respectively. In order to purify Bcr4 lacking the amino acids region 58–109 or 110–173, we amplified the DNA fragment with the primer sets of 5-bcr4-58-109-Strep-IF and 3-IF-bcr4-58-109-Strep, or 5-bcr4-110-173-Strep-IF and 3-IF-bcr4-110-173-Strep using pColdII-*bcr4*-Strep as the template. Each amplified fragment was self-ligated by the In-Fusion Cloning System, respectively, and then designated pMGKU414 or pMGKU415.

BL21 cells carrying pMGKU412, pMGKU413, pColdII-*bteA*-N-Strep (2), pMGKU414 or pMGKU415 were cultured overnight at 37°C with shaking, and then diluted 1:100 in LB liquid medium containing 50 µL/mL ampicillin and incubated for 2 h at 37°C with shaking, respectively. Each bacterial culture was further incubated overnight at 15°C in the presence of IPTG at the final concentration of 0.05 mM. Bacteria were collected by centrifugation at 2,600×*g* for 20 min, and suspended in cold TBS containing cOmplete. Each bacterial suspension was sonicated and each supernatant except for that of pMGKU413 was subjected to purification using Ni-NTA agarose (Qiagen) according to the manufacturer’s instructions. The purified proteins were dialyzed with TBS. As for pMGKU413, the bacterial suspension was sonicated and the pellet was suspended in Inclusion Body Solubilization Reagent (Thermo) according to the manufacturer’s instructions. The BscI-Strep in solubilized solution was refolded as described previously (3). Briefly, the solubilized solution was diluted with 5-fold larger refolding buffer (20 mM Tris-HCl, pH-8, 150 mM NaCl, 10% glycerol) overnight at 4°C on a rotator, and then centrifugated at 20,000×*g* for 15 min. The supernatant was dialyzed with dialysis buffer (20 mM Tris-HCl, pH-8, 150 mM NaCl, 10% glycerol), and then centrifugated at 20,000×*g* for 15 min. The supernatant was used as purified protein for the pull-down assay.

We mixed the dual-tagged protein (6×His and Strep) and 30 µl Strep-Tactin resin (IBA) in an Eppendorf tube and rotated the tube at 4°C for 1 h. Then, we washed the beads with TBS three times. Next, the V5-tagged protein-containing *E. coli* lysate was added to the tube and rotated at 4°C for 3 h. We transferred 30 µL supernatant to new Eppendorf tube and added 30 µL 2×SDS-PAGE sample buffer to prepare the Sup. Then we washed the beads with TBS (Fig. 1B and 1C) or TBS containing 0.1% Triton X-100 (Fig. 1C) three times and added 30 µL 2×SDS-PAGE sample buffer to prepare the Pellet samples.

#### Construction of a *bscI* gene-disrupted or *bspR* /*bscI* double strains

To construct the *bscI* or *bspR*/*bscI* double mutants, a 2.4 kb DNA fragment encoding *bscI* and its flanking regions was amplified by PCR with primers B1-bscI and B2-bscI using *B. bronchiceptica* S798 genomic DNA as the template. The resulting PCR product was cloned into pDONR201 to obtain pMGKU416 using the Gateway cloning system. An inverse PCR was carried out with the primers R1-bscI and R2-bscI using circular pMGKU416 as the template. The resulting PCR product was self-ligated using the In-Fusion Cloning System to obtain pMGKU417. This plasmid contained a 369-bp in-frame deletion from 30 bp downstream of the 5’ end of the *bscI* gene to 30 bp upstream of the 3’ end of the gene. This plasmid, pMGKU417, was mixed with a positive suicide vector such as pABB-CRS2 (4) to obtain pMGKU418 using the Gateway cloning system. The pMGKU418 or pABB-CRS2-*bspR* (5) plasmids in turn were introduced into *E. coli* Sm10λ*pir*, and transconjugated into the S798 wild-type or Δ*bscI* as described previously (6). The resulting mutant strains were designated Δ*bscI* or Δ*bspR*Δ*bscI*, respectively.

#### Construction of plasmids used for producing Bcr4 derivatives and BscI complementation

In order to produce full-length Bcr4 tagged with a FLAG sequence at the C-terminus, we performed an inverse PCR with primers of 5-bcr4-FLAG-IF and 3-IF-bcr4-FLAG using pDONR-*bcr4* (7) as the template. The resulting PCR product was self-ligated to obtain pMGKU419 by the In-Fusion Cloning System. To produce Bcr4 lacking the amino acid regions 159–173, 164–173 or 169–173, inverse PCR was carried out with the primer sets of 5-*bcr4*-FLAG-IF and 3-IF-bcr4Δ169-173-FLAG, 5-*bcr4*-FLAG-IF and 3-IF-*bcr4*Δ164-173-FLAG, or 5-*bcr4*-FLAG-IF and 3-IF-*bcr4*Δ159-173-FLAG using pMGKU419 as the template. Each amplified fragment was self-ligated by the In-Fusion Cloning System to obtain pMGKU420, pMGKU421 or pMGKU422, respectively. For BscI complementation, a PCR was carried out with the primers of B1-bscI-V5 and 3-B2-bscI-comp using *B. bronchiceptica* S798 genomic DNA as the template. The resulting PCR product was cloned into pDONR201 to obtain pMGKU423 using the Gateway cloning system. These plasmids were mixed with pRK-R4-R3-F, pDONR-*fha*P and pDONR-*rrnB* (8) to obtain pMGKU424, pMGKU425, pMGKU426, pMGKU427 and pMGKU428 using the Gateway cloning system.

#### Quantitative reverse transcription-PCR

Total RNA was prepared from the *B. bronchiseptica* culture using a Trizol Max Bacterial RNA isolation Kit (Invitrogen), RNeasy Mini Kit (Qiagen), and RNase-free DNase Kit (Qiagen). The reverse transcription reaction was carried out using Transcriptor Universal cDNA Master (Roche) and T100 Thermal Cycler (Bio-Rad). The quantitative RT-PCR reaction was carried out using FastStart Essential DNA Probes Master (Roche) and Light Cycler 96 (Roche). To amplify the *bscI*, *bopD*, and *recA* genes, the primer sets 5-bscI and 3-bscI, 5-bopD and 3-bopD, 5-recA and 3-recA were used, respectively. For the experiment to determine the presence of *bscI* mRNA in the Bcr4 mutant (Fig. S4), *recA* was used as an internal control. The values obtained from each strain were standardized with *recA*, and the relative amounts to the wild-type strain were determined. For the experiment to compare the amounts of *bscI* and *bopD* mRNA in the wild-type (Fig. S5), we followed the protocol provided by Roche. The genome of *B. bronchiseptica* was used as a template for quantitative RT-PCR to generate a calibration curve. The relative amount of *bopD* mRNA to *bscI* mRNA was then determined using the calibration curve.

#### Protein structure prediction

Protein structure prediction using amino acid sequences by AlphaFold2 with MMseqs2 (ColabFold) was performed on the Google Colab server with default parameters (9). A structure-based protein homology search using AlphaFold2-predicted structural models was performed on the Dali server (10). In addition, primary, secondary, and tertiary structures of query proteins were visualized and compared with those of neighbors on the Dali server.

