## Supplementary Information for "Bcr4 is a Chaperone for the Inner Rod Protein in the *Bordetella* Type III Secretion System"

BL21 cells carrying pMGKU408, pMGKU409 pMGKU410 or pMGKU411 were cultured overnight at 37ºC with shaking, and then diluted 1:40 in LB liquid medium containing 50 $\mu$L/mL ampicillin and incubated for 2 h at 37ºC with shaking. Each bacterial culture was further incubated for 5 h at 30ºC in the presence of isopropyl-beta-thiogalactopyranoside (IPTG) at the final concentration of 1 mM. Bacteria were collected by centrifugation at 2,600×*g* for 15 min, and suspended in cold TBS containing protease inhibitor cocktail, cOmplete (Roche). Each bacterial suspension was sonicated, and each supernatant was used for the pull-down assay.

Next, in order to purify Bcr4 or BscI tagged with six histidine residues (6×His) at the respective N-terminus and Strep at the respective C-terminus, we amplified DNA fragments encoding *bcr4* or *bscI* with the primer sets of 5-HindIII-bcr4 and 3-bcr4-Strep, or 5-HindIII-bscI and 3-bscI-Strep using *B. bronchiseptica* S798 genomic DNA as the template. Each amplified DNA fragment was used as a template for 2nd PCR with a primer set consisting of the upper primer used in the 1st PCR and 3-Strep-HindIII to add a 24 bp sequence encoding the Strep tag. Each resulting PCR product was cloned into the HindIII recognition sites of pColdII to obtain pMGKU412 or pMGKU413 by the In-Fusion Cloning System (Clontech), respectively. In order to purify Bcr4 lacking the amino acids region 58–109 or 110–173, we amplified the DNA fragment with the primer sets of 5-bcr4-58-109-Strep-IF and 3-IF-bcr4-58-109-Strep, or 5-bcr4-110-173-Strep-IF and 3-IF-bcr4-110-173-Strep using pColdII-*bcr4*-Strep as the template. Each amplified fragment was self-ligated by the In-Fusion Cloning System, respectively, and then designated pMGKU414 or pMGKU415.

BL21 cells carrying pMGKU412, pMGKU413, pColdII-*bteA*-N-Strep (2), pMGKU414 or pMGKU415 were cultured overnight at 37ºC with shaking, and then diluted 1:100 in LB liquid medium containing 50 $\mu$L/mL ampicillin and incubated for 2 h at 37ºC with shaking, respectively. Each bacterial culture was further incubated overnight at 15ºC in the presence of IPTG at the final concentration of 0.05 mM. Bacteria were collected by centrifugation at

2,600×*g* for 20 min, and suspended in cold TBS containing cOmplete. Each bacterial suspension was sonicated and each supernatant except for that of pMGKU413 was subjected to purification using Ni-NTA agarose (Qiagen) according to the manufacturer's instructions. The purified proteins were dialyzed with TBS. As for pMGKU413, the bacterial suspension was sonicated and the pellet was suspended in Inclusion Body Solubilization Reagent (Thermo) according to the manufacturer's instructions. The BscI-Strep in solubilized solution was refolded as described previously (3). Briefly, the solubilized solution was diluted with 5-fold larger refolding buffer (20 mM Tris-HCl, pH-8, 150 mM NaCl, 10% glycerol) overnight at 4°C on a rotator, and then centrifugated at 20,000×*g* for 15 min. The supernatant was dialyzed with dialysis buffer (20 mM Tris-HCl, pH-8, 150 mM NaCl, 10% glycerol), and then centrifugated at 20,000×*g* for 15 min. The supernatant was used as purified protein for the pull-down assay.

We mixed the dual-tagged protein (6×His and Strep) and 30 *μ*l Strep-Tactin resin (IBA) in an Eppendorf tube and rotated the tube at 4°C for 1 h. Then, we washed the beads with TBS three times. Next, the V5-tagged protein-containing *E. coli* lysate was added to the tube and rotated at 4°C for 3 h. We transferred 30 *μ*L supernatant to new Eppendorf tube and added 30 *μ*L 2×SDS-PAGE sample buffer to prepare the Sup. Then we washed the beads with TBS (Fig. 1B and 1C) or TBS containing 0.1% Triton X-100 (Fig. 1C) three times and added 30 *μ*L 2×SDS-PAGE sample buffer to prepare the Pellet samples.

Table S1. Nomencalture of the *Bordetella* T3SS

| Unified name | *Bordetella* | *Yersinia* | *Pseudomonas* | Predicted function |
|---|---|---|---|---|
| SctF | BscF | YscF | PscF | Needle |
| SctI | BscI | YscI | PscI | Inner rod |
| SctE | BopB | YopB | PopB | Translocation pore |
| SctB | BopD | YopD | PopD | Translocation pore |
| SctA | Bsp22 | LcrV | PcrV | Needle tip or Filament |
| SctJ | BscJ | YscJ | PscJ | Inner membrane ring |
| SctK | BscK | YscK | PscK | ATPase cofactor |
| SctN | BscN | YscN | PscN | ATPase |

| Table S2. Plasmids used in the study | | |
|---|---|---|
| Name | Description | Reference or source |
| pDONR201 | DNA cloning vector, Km$^r$ | Invitrogen |
| pMGKU404 | pDONR201, *bscI* gene tagged with V5 sequence at the C terminus | This study |
| pMGKU405 | pDONR201, *bscK* gene tagged with V5 sequence at the C terminus | This study |
| pMGKU406 | pDONR201, *bscJ* gene tagged with V5 sequence at the C terminus | This study |
| pMGKU407 | pDONR201, *bcr4* gene tagged with V5 sequence at the C terminus | This study |
| p99*ccdB*-V5 | Expression vector for V5 tagged gene, Amp$^r$ | 1 |
| pMGKU408 | p99-*ccdB*-V5, *bscI* gene tagged with V5 sequence at the C terminus | This study |
| pMGKU409 | p99-*ccdB*-V5, *bscK* gene tagged with V5 sequence at the C terminus | This study |
| pMGKU410 | p99-*ccdB*-V5, *bscJ* gene tagged with V5 sequence at the C terminus | This study |
| pMGKU411 | p99-*ccdB*-V5, *bsc4* gene tagged with V5 sequence at the C terminus | This study |
| pColdII | Expression vector for His tagged gene, Amp$^r$ | TAKARA |
| pMGKU412 | pColdII, *bcr4* gene tagged with Strep sequence at the C terminus | This study |
| pMGKU413 | pColdII, *bscI* gene tagged with Strep sequence at the C terminus | This study |
| pMGKU414 | pColdII, *bcr4* gene lacking amino acids region 58-109 tagged with Strep sequence at the C terminus | This study |
| pMGKU415 | pColdII, *bcr4* gene lacking amino acids region 110-173 tagged with Strep sequence at the C terminus | This study |
| pColdII-*bteA*-N-Strep | pColdII, *bteA* gene coding amino acids region 1-312 tagged with Strep sequence at the C terminus | 2 |
| pMGKU416 | pDONR201, *bscI* gene | This study |
| pMGKU417 | pDONR201, *bscI* gene containing internal sequence-deletion and its flanking region | This study |
| pABB-CRS2 | Suicide vector for conjugation, Amp$^r$ | 4 |
| pMGKU418 | pABB-CRS2, *bscI* gene containing internal sequence-deletion and its flanking region | This study |
| pABB-CRS2-*bspR* | pABB-CRS2, *bspR* gene containing internal sequence-deletion and its flanking region | 5 |
| pDONR-*bcr4* | pDONR201, *bcr4* gene for complementation | 7 |
| pMGKU419 | pDONR201, *bcr4* gene tagged with FLAG sequence at the C terminus for complementation | This study |
| pMGKU420 | pDONR201, *bcr4* gene lacking amino acids region 169-173 tagged with FLAG sequence at the C terminus | This study |
| pMGKU421 | pDONR201, *bcr4* gene lacking amino acids region 164-173 tagged with FLAG sequence at the C terminus | This study |
| pMGKU422 | pDONR201, *bcr4* gene lacking amino acids region 159-173 tagged with FLAG sequence at the C terminus | This study |
| pMGKU423 | pDONR201, *bscI* gene for complementation | This study |
| pRK415-R4-R3-F | pRK415, recombination sites for MultiSite Gateway, Tet$^r$ | 8 |
| pDONR-*fhaP* | pDONR-P4-P1R, *fha* promoter | 8 |
| pDONR-*rrnB* | pDONR-P2R-P3, *rrnB* terminator | 8 |
| pMGKU424 | pRK415-R4-R3-F, *bcr4* gene tagged with FLAG sequence at the C terminus for complementation | This study |
| pMGKU425 | pRK415-R4-R3-F, *bcr4* gene lacking amino acids region 169-173 tagged with FLAG sequence at the C terminus | This study |
| pMGKU426 | pRK415-R4-R3-F, *bcr4* gene lacking amino acids region 164-173 tagged with FLAG sequence at the C terminus | This study |
| pMGKU427 | pRK415-R4-R3-F, *bcr4* gene lacking amino acids region 159-173 tagged with FLAG sequence at the C terminus | This study |
| pMGKU428 | pRK415-R4-R3-F, *bscI* gene | This study |

| Table S3. Primers used in the study | |
|---|---|
| Name | Sequence (5'→3') |
| B1-bscI-V5 | AAAAAGCAGGCTTGTTGGACAGCGGACCCGCC |
| B2-bscI-V5 | AGAAAGCTGGGTTTGACATTCTCGCCAGCGTGTC |
| B1-bscK-V5 | AAAAAGCAGGCTTGGCGCACGCGCTGGTCCCG |
| B2-bscK-V5 | AGAAAGCTGGGTTGGCTTGGGCGGGGAACGAGG |
| B1-BscJ-V5 | AAAAAGCAGGCTTGGAGTACGAGTTGGTGGGC |
| B2-BscJ-V5 | AGAAAGCTGGGTTGTCATGCCCGGCTCCTTCCG |
| B1-bcr4-comp | AAAAAGCAGGCTGCCAGGTCCGGTCTCGCACCG |
| B2-bcr4-V5 | AGAAAGCTGGGTTTCCAGGAGCTCCAGGTAATG |
| 5-HindIII-bcr4 | ATCCGAATTCAAGCTTCATTCAGACTCAGGTTCAGATTC |
| 3-bcr4-Strep | CGGGTGGCTCCATCCAGGAGCTCCAGGTAATG |
| 5-HindIII-bscI | ATCCGAATTCAAGCTTAATTTGGATCTGACGGCGATC |
| 3-bscI-Strep | CGGGTGGCTCCATGACATTCTCGCCAGCGTGTC |
| 3-Strep-HindIII | GCAGGTCGACAAGCTTTCATTTTTCGAACTGCGGGTGGCTCCA |
| 5-bcr4-58-109-Strep-IF | GCCGTCAGCGATCTGTCGGACTGGACGCCTGGTG |
| 3-IF-bcr4-58-109-Strep | CAGATCGCTGACGGCCGCCG |
| 5-bcr4-110-173-Strep-IF | TGGAGCCACCCGCAGTTCGA |
| 3-IF-bcr4-110-173-Strep | CTGCGGGTGGCTCCAGCGGTCCTCCGCCTGTGCCG |
| B1-bscI | AAAAAGCAGGCTTCATGACTGTTCACGACGAC |
| B2-bscI | AGAAAGCTGGGTGCACAAGATCCAGCGCGACC |
| R1-bscI | CGACGTTGGCGTTGATCGCCGTCAGATC |
| R2-bscI | TCAACGCCAACGTCGACACGCTGGCGAG |
| 5-bcr4-FLAG-IF | CCACCCGCAGTTCGAAAAATGAATTTGGATCTACCCAGCTTTCTTG |
| 3-IF-bcr4-FLAG | TCGAACTGCGGGTGGCTCCATCCAGGACCTCCAGGTAATGGCTCG |
| 3-IF-bcr4∆169-173-FLAG | GTCATCCTTGTAGTCTAATGGCTCGGACTGCAACG |
| 3-IF-bcr4∆164-173-FLAG | GTCATCCTTGTAGTCCAACGGCGCGGGCCGCATCATG |
| 3-IF-bcr4∆159-173-FLAG | GTCATCCTTGTAGTCCATCATGGCGGGTCCGCTGTC |
| 3-B2-bscI-comp | AGAAAGCTGGGTTTGACATTCTCGCCAGCGTGTC |
| 5-bscI | CGATCTGCAGGCCAGGTT |
| 3-bscI | CTATTGCCTTGCCCACCAA |
| 5-bopD | CGGCTCGGTGAAGACATC |
| 3-bopD | CCTCCCGCATCTGTTGAC |
| 5-recA | ATGAAGATCGGCGTGATGT |
| 3-recA | TAGAACTTGAGCGCGTTGC |

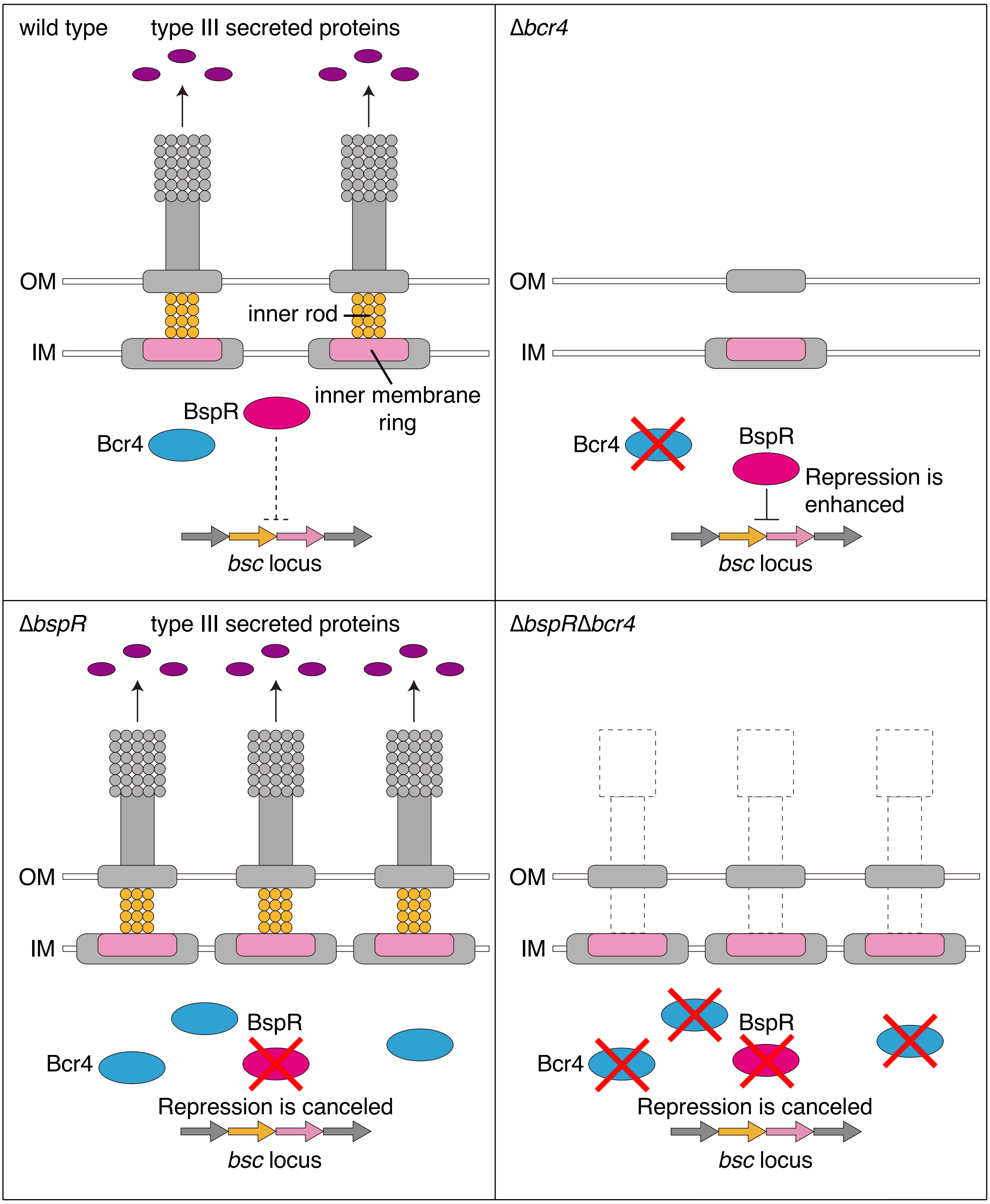


**Fig. S1. Construction of the T3SS machinery in strains lacking Bcr4 and/or BspR.** In the *B. bronchiseptica* wild-type (upper left), the BspR negative regulation level for the *bsc* locus transcription is moderate, and the T3SS machinery is established. In the Bcr4-deficient strain (upper right), BspR strongly represses the *bsc* locus transcription, and construction of the T3SS machinery is incomplete. In the BspR-deficient strain (lower left), the negative regulatory effect of BspR is cancelled, and the construction of the T3SS machinery is promoted. In the BspR/Bcr4 double-deficient strain (lower right), while the *bsc* locus transcription is promoted because of BspR deficiency, T3SS is not functional.

Fig. S1

```
Bb       1 MHSDSGSDSGSDSGSGS--PMASSIHPSEPIQPMEHVLEEADARLLTEVGFLAAAVSDLT     58
Bp       1 MHSDSGSDSGSDSGSGS--PMVSSIHPSEPIQPMEHVLEEADARLLTEVGFLAAAVSDLT     58
Bpp      1 MHSDSGSDSGSGSGSGSGSPMASSIHPSEPIQPMEHVLEEADARLLTEVGFLAAAVSDLT     60

Bb      59 RADAIFNALQRVRPGRTYPCIGLAVARMNAGLPDEAAEILANFQPAQAEDRSELDAWCGF    118
Bp      59 RADAIFNALQRVRPGRTHPCIGLAVARMNAGLPDEAAEILANFQPAQPEDRSELDAWCGF    118
Bpp     61 RADAIFNALQRVRPGRTYPCIGLAVARMNAGLPDEAAEVLANFQPAQAEDRSELDAWCGF    120

Bb     119 ALLLAGRSDEARRMLQRAIDAGGEAARLAQVVLDSGPAMMRPAPLQSEPLPGAPG         173
Bp     119 ALLLAGRSDEARRMLQRAIDAGGEAARLAQVVLDSGPAMMRPAPLQSEPLPGAPG         173
Bpp    121 ALLLAGRSDEARRMLQRAIDAGGEAARLAQVVLDSGPAMMRPAPLQSEPLPGAPG         175
```

**Fig. S2. Alignment of Bcr4 amino acid sequences in representative *Bordetella* species.** Bcr4 amino acid sequences of *B. bronchiseptica* S798 (Bb), *B. pertussis* Tohama I (Bp) and *B. parapertussis* 12822 (Bp) were compared using ClustalW. The grey highlighted letters represent amino acid residues that were different from those in Bb. Bcr4 of Bp and Bpp have 98.3% and 97.1% identities with those of Bp, respectively.

Fig. S2

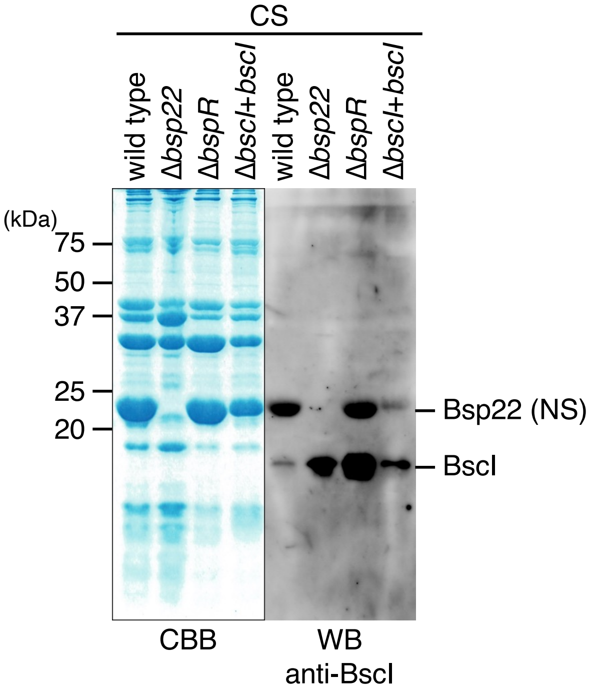


**Fig. S3. The nonspecific reaction of anti-BscI antibody to Bsp22.** The culture supernatants (CS) were prepared from the wild-type strain, Δ*bsp22* (Bsp22-deficient strain), Δ*bspR* (BspR-deficient strain) or Δ*bscI*+*bscI* (BscI-complemented strain) cultured in SS medium. The CS samples were separated by SDS-PAGE and stained with Coomassie Brilliant Blue (CBB, left panel) or analyzed by Western blotting (WB) with anti-BscI antibody (right panel). NS indicates nonspecific signals. Experiments were performed at least three times, and representative data are shown.

Fig. S3

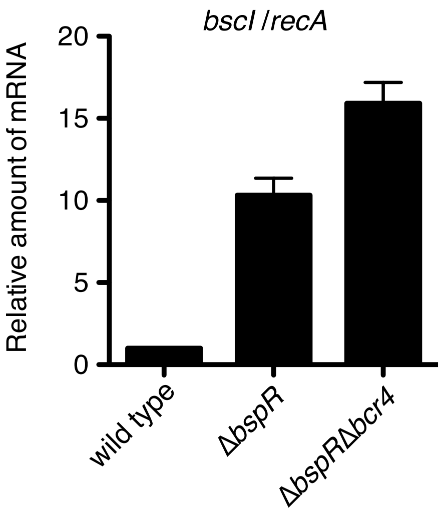


**Fig. S4. The results of the RT-PCR analysis for the mRNA level of *bscI* in *B. bronchiseptica* strains.** Total RNA was prepared from the wild-type strain, Δ*bspR* (BspR-deficient strain) or Δ*bspR*Δ*bcr4* (BspR- and Bcr4-deficient strain) cultured in SS medium and subjected to a quantitative RT-PCR analysis. The histogram shows the relative amount of *bscI* mRNA normalized by the housekeeping gene, *recA* mRNA. Experiments were performed at least three times, and representative data are shown.

Fig. S4

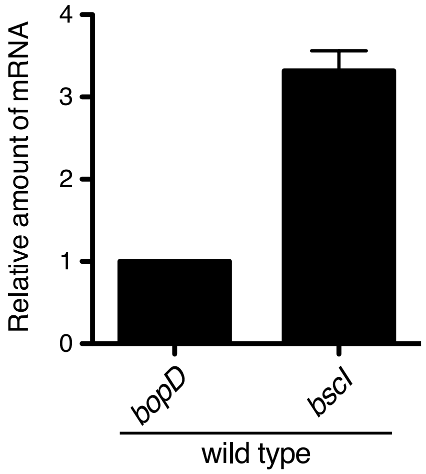


**Fig. S5. The results of the RT-PCR analysis for mRNA levels of *bopD* and *bscI* in the wild-type *B. bronchiseptica*.** Total RNA was prepared from the wild-type strain cultured in SS medium and subjected to a quantitative RT-PCR analysis. The histogram shows the relative amount of *bopD* and *bscI* mRNA in the wild-type. The relative ratio of *bscI* mRNA is shown when the *bopD* mRNA amount is set as 1. Experiments were performed at least three times, and representative data are shown.

Fig. S5

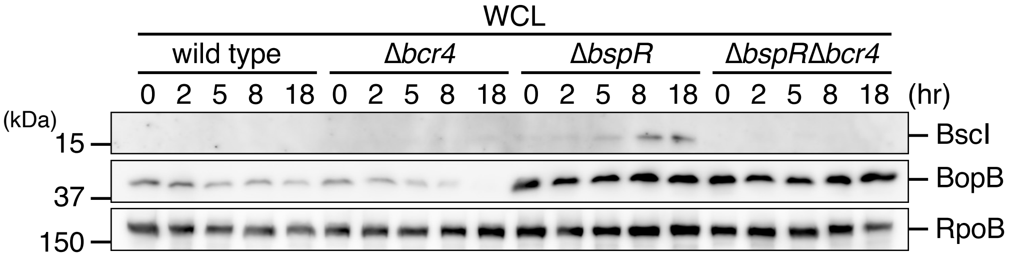


**Fig. S6. The time course of BscI production in *B. bronchiseptica*.** The whole cell lysates (WCL) were prepared from the wild-type strain, Δ*bcr4* (Bcr4-deficient strain), Δ*bspR* (BspR-deficient strain) or Δ*bspR*Δ*bcr4* (BspR- and Bcr4-deficient strain) cultured in SS medium for 0, 2, 5, 8 or 18 hr. The WCL were separated by SDS-PAGE and analyzed by Western blotting with antibodies against BscI, BopB and RpoB. Experiments were performed at least three times, and representative data are shown.

Fig. S6

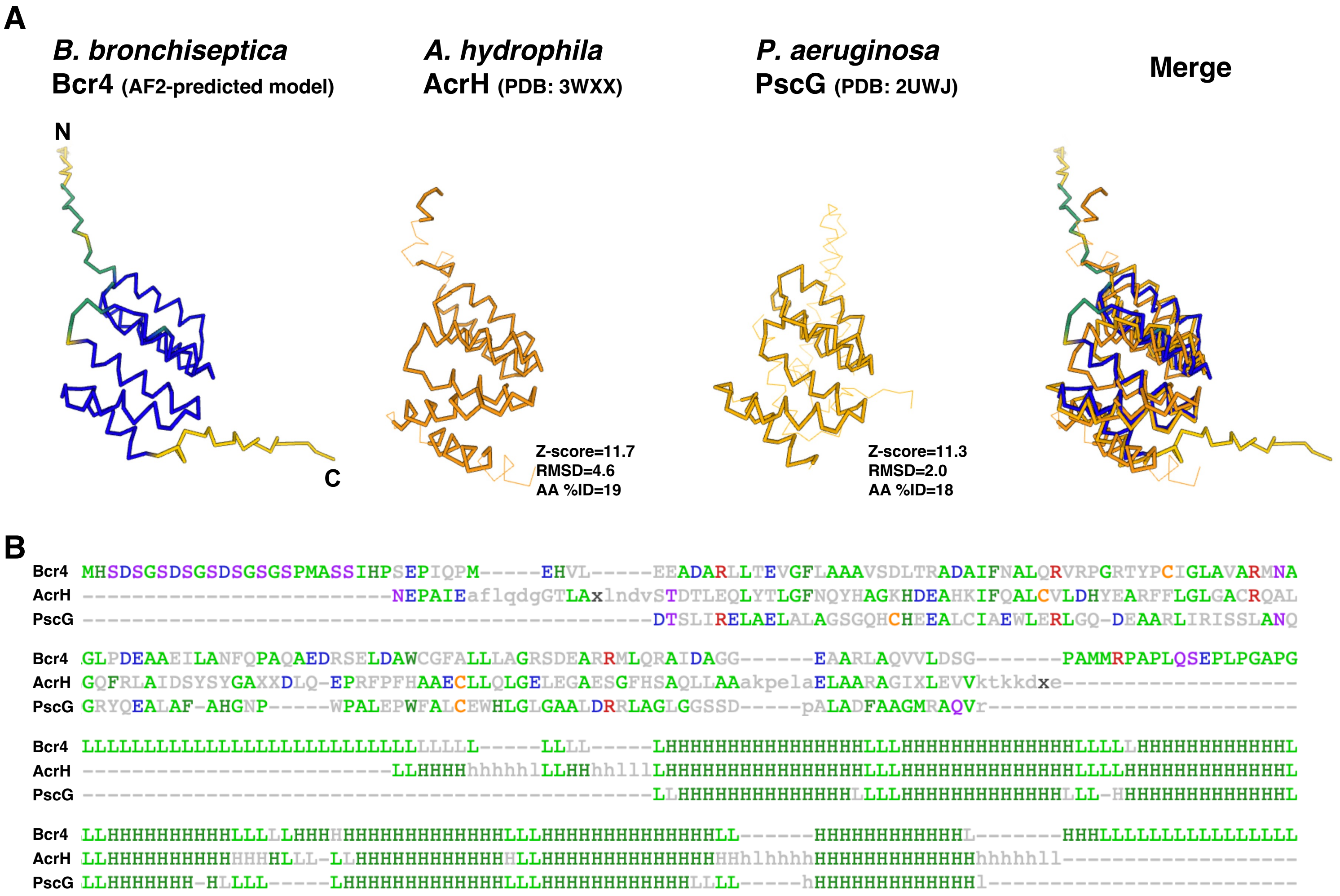


**Fig. S7. The predicted structural model of *B. bronchiseptica* Bcr4**

(A) The AlphaFold2 (AF2)-predicted structural model of *B. bronchiseptica* Bcr4 and structural comparison with *Aeromonas hydrophil*a AcrH (PDB: 3WXX) and *Pseudomonas aeruginosa* PscG (PDB: 2UWJ). Z-score, root mean square deviation (RMSD), and amino acid identity (AA %ID) of AcrH and PscG compared with Bcr4 are shown, respectively. (B) The pairwise sequence alignment of Bcr4, AcrH, and PscG. The most frequent amino acid type is colored. The secondary structure assignments (H/h: helix, E/e: strand, L/l: coil) are also shown.

Fig. S7
